# Mef-2 and p300 interact to regulate expression of the homeostatic regulator Pumilio in *Drosophila*

**DOI:** 10.1101/264911

**Authors:** Wei-Hsiang Lin, Richard A. Baines

## Abstract

Pumilio (Pum) is a key component of neuron firing-rate homeostasis that maintains stability of neural circuit activity. Whilst Pum is ubiquitously expressed, we understand little about how synaptic excitation regulates its expression. Here, we characterised the *Drosophila dpum* promoter and identified multiple Myocyte enhancer factor-2 (Mef2)-binding elements. To understand the transactivation capability of dMef2, we cloned 12 *dmef2* splice variants and used a luciferase-based assay to monitor *dpum* promoter activity. Whilst all 12 dMef2 splice variants enhance *dpum* promoter activity, exon 10-containing variants induce greater transactivation. Previous work shows dPum expression increases with synaptic excitation. However, we observe no change in *dmef2* transcript in CNS exposed to picrotoxin (PTX). The lack of activity-dependence is indicative of additional regulation. We identified p300 as a likely candidate. We show that by binding to dMef2, p300 represses *dpum* transactivation. Significantly, *p300* transcript is down-regulated by enhanced synaptic excitation (PTX) which, in turn, increases transcription of *dpum* through derepression of dMef2. These results suggest the activity-dependent expression of *dpum* is regulated by an interaction between p300 and dMef2.

## Introduction

Pumilio (Pum), a founding member of the Pum/FBF (Puf) RNA-binding protein family, is central to multiple aspects of CNS function, including (but not limited to) firing-rate homeostasis, dendritic morphogenesis, synaptic growth and function, expression of acetylcholinesterase and long-term memory (Chen et al., 2008, Driscoll et al., 2013, Menon et al., 2004, Muraro et al., 2008, Vessey et al., 2010). Despite a wide-ranging involvement in many aspects of CNS function, little is understood concerning the regulation of Pum expression in the CNS. Importantly, reduced levels of Pum have been linked to epilepsy in *Drosophila*, rodents and human (Follwaczny et al., 2017, Lin et al., 2017, Siemen et al., 2011, Wu et al., 2015).

Pum binds an 8 nucleotide sequence in mRNA (UGUANAUA, where N = A, G, C or U), termed a Pum Response Element (PRE) and, by doing so, induces translational repression (Arvola et al., 2017, Wharton et al., 1998, Wreden et al., 1997). Pum-dependent translational repression requires a number of co-regulators, including Nanos (Nos) and Brain tumor (Brat), which bind different, but characterised, RNA motifs to form a complex with Pum (Arvola et al., 2017). An analysis of 3’UTRs in the *Drosophila* genome identified 2477 transcripts containing one or more PREs highlighting the possibility that many transcripts undergo Pum-mediated translational regulation. The number of transcripts regulated may, however, be considerably less because specificity is also likely provided by both PRE copy-number and proximity of PRE-, Nos- and Brat-binding motifs within individual transcripts (Arvola et al., 2017).

The number of transcripts expressing PREs is indicative of the importance of Pum. Despite this, however, our understanding of *pum* expression and role(s) is limited and, where information is known, is mostly focused on post-transcriptional modification. For example, the *dpum* transcript is itself regulated through translational repression by the cytoplasmic RNA-Binding Fox protein (Rbfox1, aka A2BP1) in order to promote germ cell development (Carreira-Rosario et al., 2016). In mammals, Myocyte Enhancer Factor-2 (Mef2) regulates expression of miR-134 which, in turn, downregulates *pum2* transcript to fine tune dendrite morphogenesis (Fiore et al., 2009, Fiore et al., 2014). Mef2 is an activity-dependent transcription factor that has been implicated to control synapse formation in addition to dendrite morphogenesis (Flavell et al., 2006). Depending on interaction with either positive or negative cofactors, Mef2 can potentiate or repress gene transcription. Through an interaction with GATA4, a cardiac-enriched transcription factor, Mef2 activates the *Nppa* promoter to regulate cardiac development (Morin et al., 2000). In contrast, Mef2 forms a complex with class II histone deacetylases (HDACs) to repress gene transcription by deacetylating histones, resulting in chromatin condensation and a reduced accessibility of core transcriptional machinery to promoter region of target genes (Kao et al., 2001, Lu et al., 2000, McKinsey et al., 2001).

To identify how transcription of *pum* is regulated, we cloned the promoter region of *dpum* and identified putative binding motifs for 114 transcription factors, including multiple dMef2 elements. A luciferase-based reporter, driven by the *dpum* promoter, shows that dMef2 is sufficient to transactivate the *dpum* promoter. Magnitude of transactivation varies across the many dMef2 splice-variants present in CNS. Significantly, we also report that dMef2 mediated transactivation of *dpum* is repressed by p300 (aka Nejire), a histone acetyltransferase (HAT). Unlike dMef2, we show that *p300* expression is directly regulated by neuronal activity and, thus, provide a potential route through which membrane depolarization regulates expression level of *dpum*.

## Results

### Identification of the *pumilio* promoter region

To understand transcriptional regulation of *dpum*, we identified the *dpum* promoter region. A 2-kb region upstream of the transcription start site was targeted as a potential location. Interrogation of transcription factor databases (TRANSFAC model, MAPPER) (Marinescu et al., 2005), identified putative binding motifs for 114 transcription factors within the region -2000 to +1 (transcription initiation marked as +1, motifs listed in Table EV1). Putative transcriptional binding motifs include: Sp1, TBP, C/EBP, Oct-1, Mef2, MADS-A/B, Hb, NF-kappaB, TCF, CREB and SRF (Fig. EV1). To test for function of the *dpum* promoter region, the 2-kb fragment, (-2000 to +1, termed *pumA*) was placed upstream of *firefly*-luciferase (FF) and transiently transfected in S2R+ cells. After 24 h *pumA:FF* resulted in a 156.5 ± 21.9-fold increase in FF activity compared to transfection of cells with empty vector (set at 1, Fig. 1, *p* = 2.5×10^-7^).

**Figure 1.**
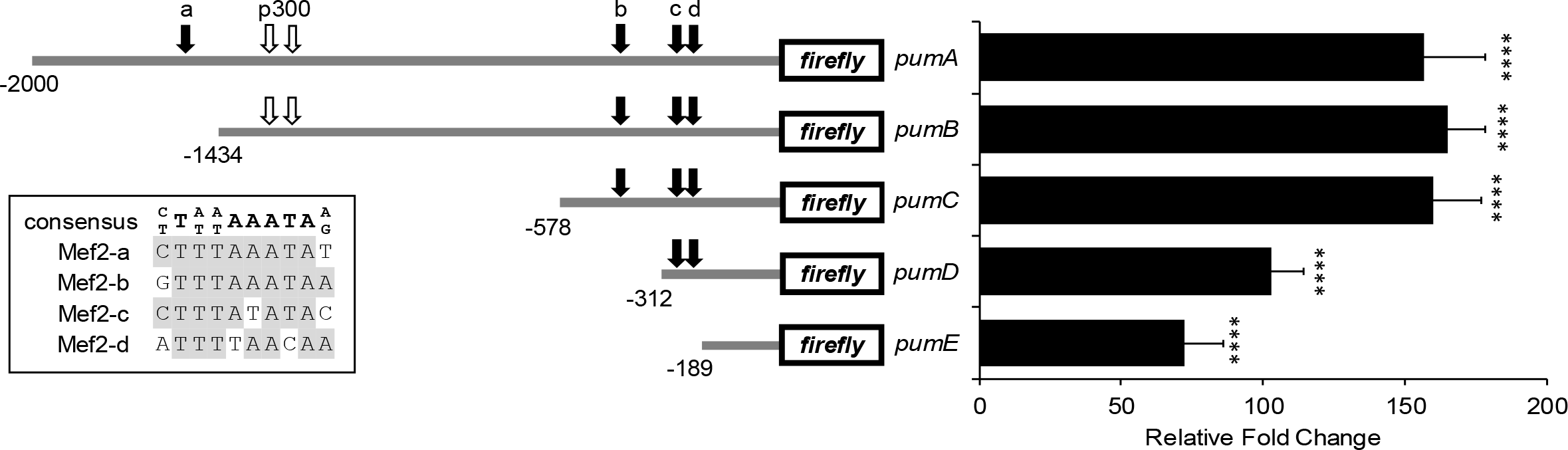
Characterisation of *dpum* promoter activity. Activity analysis, by luciferase assays, of constructs bearing defined regions of a putative 2-kb *dpum* promoter that was placed upstream offirefy-luciferase (FF). These regions are termed *pumA, pumB, pumC, pumD* and *pumE*, respectively (-2000, -1434, -578, -312 and -189 to +1: transcription initiation marked as +1). Constructs were transiently co-transfected, together with a *renilla*-luciferase (*Ren*, a loading control, driven by *actin* promoter), in S2R+ cells for 24 h and *pum:FF* to *Ren* ratio was calculated. *PumA, pumB, pumC, pumD* and*pumE:FF* resulted in 156.5 ± 21.9, 164.8 ± 13.7, 159.8 ± 17.2, 103 ± 11.1 and 72.5 ± 13.4-fold increase in FF activity compared to transfection of cells with empty vector (FF to Ren ratio was set at 1) (n = 5 independent transfections). Four putative Mef2 binding sites (Mef2-a, -b, -c and -d) with a consensus sequence (C/T)T(A/T)(A/T)AAATA(A/G) were identified (black arrows) and each binding sequence is shown in the inset (identical nucleotides shown in grey boxes). White arrows indicate two potential p300 binding sites. Data information: Data are presented as mean ± s.d. ****P ≤ 0.0001 (ANOVA with Bonferroni’s post-hoc).

To evaluate the minimal promoter region of *dpum*, we generated a series of deletion constructs in addition to *pumA: pumB* (−1434 to +1), *pumC* (-578 to +1), *pumD* (-312 to +1) and *pumE* (-189 to +1) (Fig. 1). Compared to *pumA:FF*, *pumB:FF* and *pumC:FF* resulted in a similar increase in FF expression (164.8 ± 13.7 and 159.8 ± 17.2-fold, *p* = 4×10^-9^ and 3.2×10^-8^, respectively), whilst *pumD:FF* and *pumE:FF* showed a lower, but still significant, increase (103 ± 11.1 and 72.5 ± 13.4-fold increase, *p* = 3.2×10^-8^ and 2.2×10^-6^, respectively) (Fig. 1). Thus, it would seem that multiple elements contained within the 2-kb region are capable of increasing *dpum* expression.

### dMef2 transactivates the *pumilio* promoter

Mef2 is reported to reduce *pum2* transcript post-transcriptionally through increased expression of miR-134 (Fiore et al., 2009). However, our analysis of the 2-kb *dpum* promoter identified four putative Mef2 binding sites with the consensus sequence (C/T)T(A/T)(A/T)AAATA(A/G) (Gossett et al., 1989) (Fig. 1). These four Mef2 elements (termed Mef2-a, -b, -c and -d), are located at -1561, -423, -298 and -214bp, respectively (Fig. 1). This high number of sites is indicative of direct regulatory effect. To test if *dpum* expression is regulated by Mef2, we used *Drosophila mef2 (dmef2)* dsRNA (specific for all variants) to knockdown endogenous expression in S2R+ cells that co-expressed the *pumA:FF* reporter. This was sufficient to reduce *pumA* promoter activity (154.5 ± 13.1 vs. 113.2 ± 15.1 control vs. *dmef2* dsRNA, *p* = 0.006) indicative that *dpum* expression is endogenously regulated by dMef2.

*Dmef2* is encoded by a single gene (Lilly et al., 1994, Nguyen et al., 1994, Taylor et al., 1995) and contains 15 exons. Exons *10* and *14* are alternatively spliced, while exons *9* and *15* contain cryptic splice sites and generate cassettes 9A and 15A, respectively (Fig. 2A). To determine the most common splice isoforms of *dmef2* transcripts present in CNS of 3^rd^ instar *Drosophila* larvae, RT-PCR was used to isolate and clone 56 complete open reading frames (ORFs). Comparison of exon composition of clones revealed 9 unique splice variants. Isoforms *dmef2(I-IV)* were previously identified (Gunthorpe et al., 1999, Taylor et al., 1995), whilst *dmef2(V-VIII)* and *dmef2(mini)* are novel. Analysis of the 56 ORFs showed that *dmef2(II)* and *dmef2(VI)* are present at highest frequency (Fig. 2A). Analysis of exon usage across all splice sites show that exon *10* is present at highest abundance followed by exon *14* (Fig. 2B).

**Figure 2.**
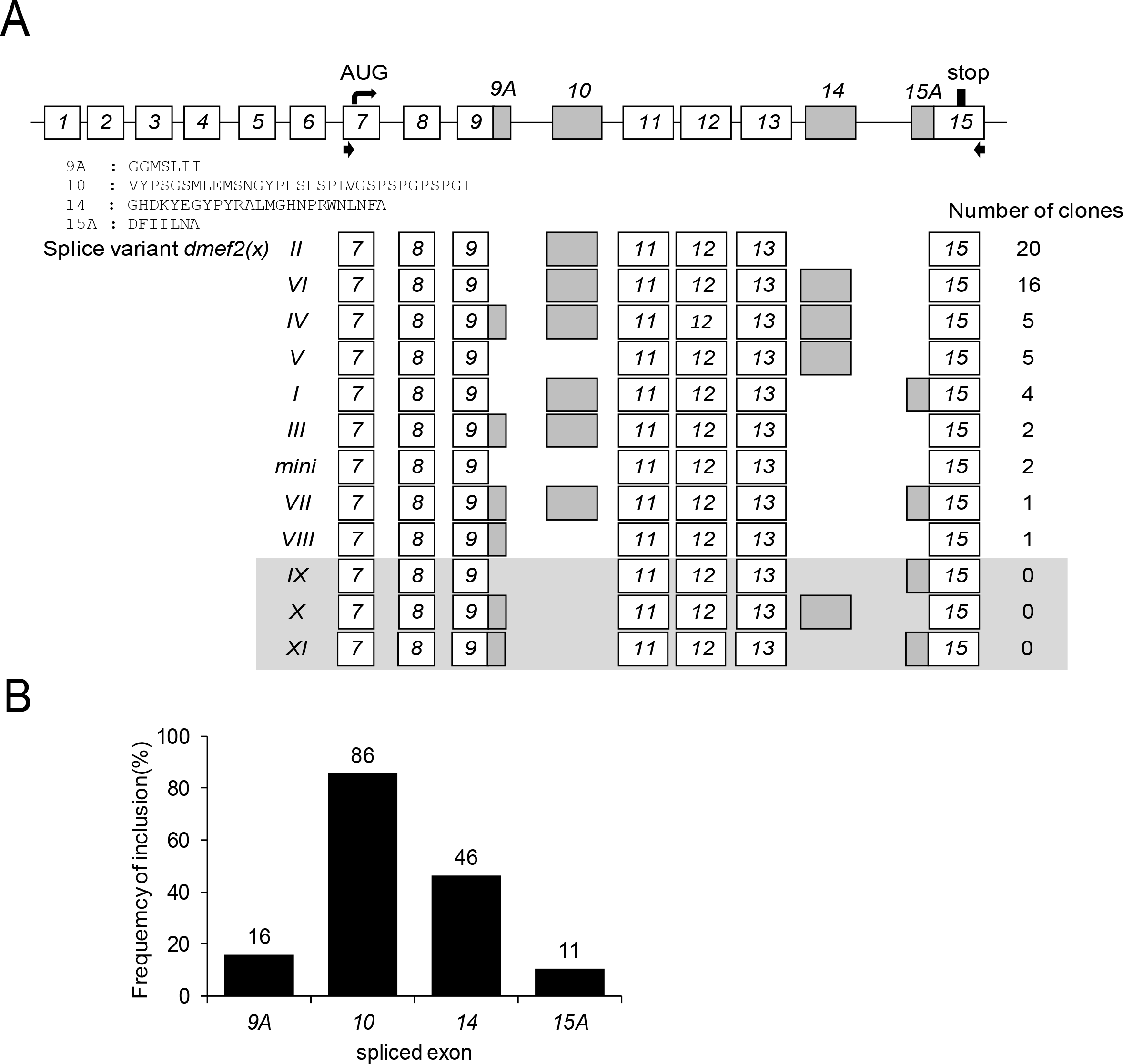
Characterisation of splice variants of *dmef2* isolated from 3^rd^ instar larval central nervous system. A Schematic of the *dmef2* gene structure. *Dmef2* contains 15 exons with exon *10* and *14* being alternatively spliced, while exons *9* and *15* contain cryptic splice sites and generate cassettes 9A and 15A, respectively. The amino acid sequences of exons 9A, 10, 14 and 15A are shown. Black arrows indicate the location of primer pairs used to amplify the open reading frame of *dmef2.* Exon usage of splice variants, termed *dmef2(I-VIII)* and *dmef2(mini)*, and the frequency of clones are indicated. The clones, *dmef2(IX-XI)*, in the shaded box were not found in the 3^rd^ instar CNS, but are theoretically possible. B Analysis of exon usage across the identified *dmef2* splice variants shows that exon *10* is present at highest abundance (86%) followed by exon *14* (46%).

To have a better understanding of the ability of individual dMef2 splice variants to transactivate *pumA:FF* activity, we constructed another 3 splice variants (although not found in our CNS analysis, these variants are theoretically possible), *dmef2(IX-XI)* (Fig. 2A). Expression of individual *dmef2* variants, in S2R+ cells, resulted in a 1.6 ± 0.1 to 3.3 ± 0.2-fold increase in FF expression compared to control (no *dmef2* expression, set at 1) (Fig. 3A). Sorting splice variants by fold change formed a clear cluster containing exon *10* (Fig. 3B). Exon *10* is contained within *dmef2(I, II, III, IV, VI* and *VII*) that, collectively, transactivated the *dpum* promoter to a greater level than variants lacking this exon: *dmef2(V, VIII, IX, X, XI* and *mini*). Comparing identical *dmef2* variants that differ only by the inclusion of exon *10* (e.g. *dmef2(I)* vs. *(IX); (II)* vs. *(mini); (III)* vs. *(VIII); (IV)* vs. *(X)* and *(VII)* vs. *(XI))* confirms that inclusion of this exon results in significant transactivation.

**Figure 3.**
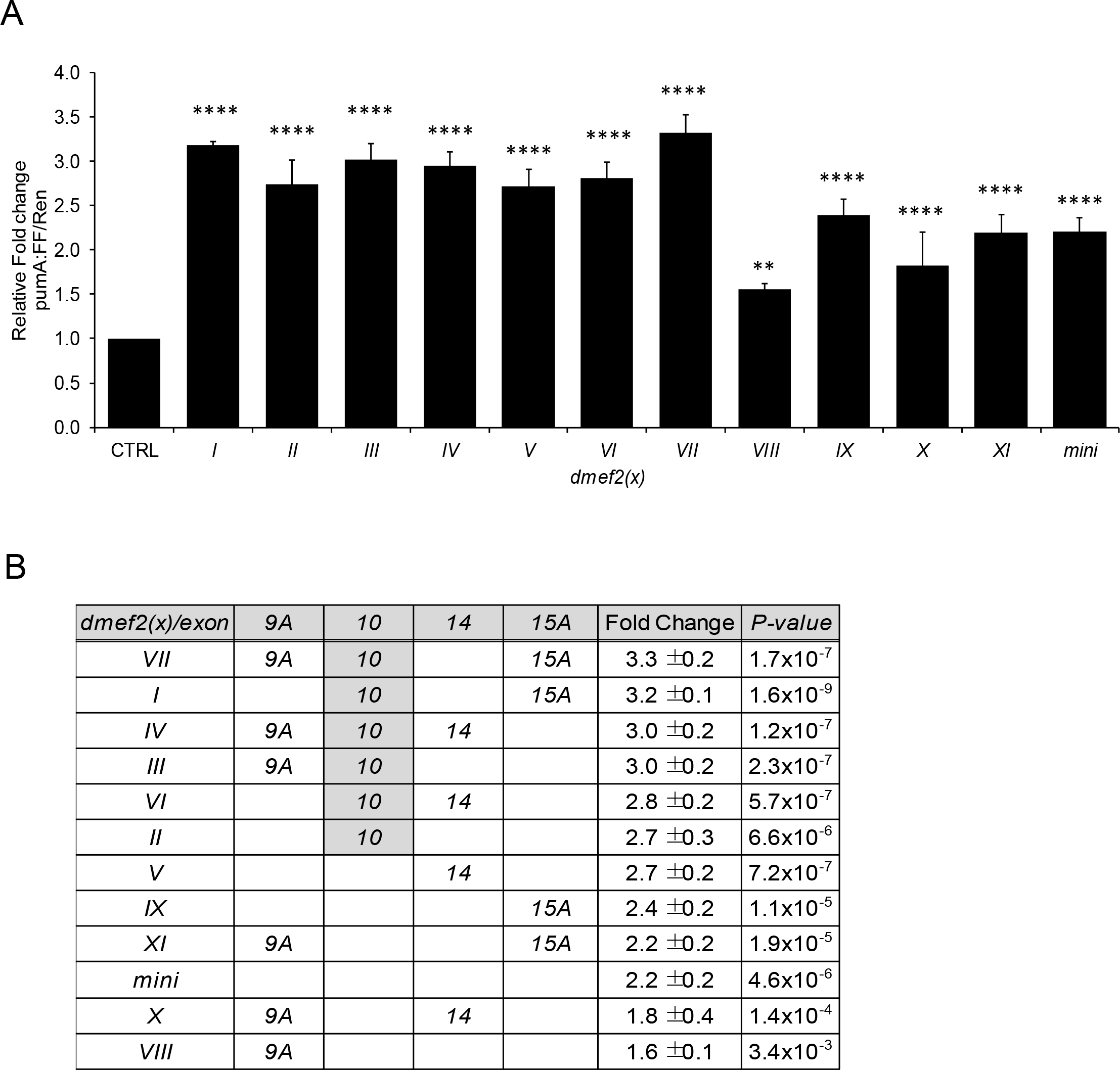
dMef2 splicing variants transactivate the *dpum* promoter. A Shows expression of *pumA* promoterfirefy-luciferase *(pumA:FF)* reporter with individual *dmef2* splice variants *(dmef2(x))* and a renilla-luciferase gene *(Ren*, loading control, driven by *actin* promoter) in S2R+ cells. Expression of *dmef2* variants resulted in a 1.6 ± 0.1 to 3.3 ± 0.2-fold increase in luciferase activity compared to control (CTRL, no *dmef2* expression, set at 1) (n = 5 independent transfections). B Shows the effectiveness of each splice variant ranked highest to lowest. Exon *10* containing variants, *dmef2(I, II, III, IV, VI and VII)*, transactivated the *dpum* promoter to a greater level than variants lacking this exon: *dmef2(V, VIII, IX, X, XIand mini).* Data information: Data are presented as mean ± s.d. **P ≤ 0.01 and ****P ≤ 0.0001 (ANOVA with Bonferroni’s post-hoc).

To further confirm dMef2 transactivation of *dpum*, we compared FF activity of each *dpum* promoter construct *(pumA* to *E)* following *dmef2(VII)* overexpression (the variant showing the strongest transactivation). Promoter fragments *pumA-E* contain 4, 3, 3, 2 and 0 predicted Mef2 binding elements, respectively (Fig. 1). Overexpression of *dmef2* resulted in 1.9 ± 0.3, 1.7 ± 0.1, 1.4 ± 0.3 and 1.4 ± 0.3-fold increase in promoter activity *(pumA-D, p* = 1.2×10^-6^, 4.1×10^-6^, 0.004, 0.05 respectively) compared to control (no *dmef2* expression, set at 1), whilst *pumE* (lacking an Mef2 binding motif) showed no change (1.2 ± 0.1, *p* > 0.05) (Fig. 4A).

**Figure 4.**
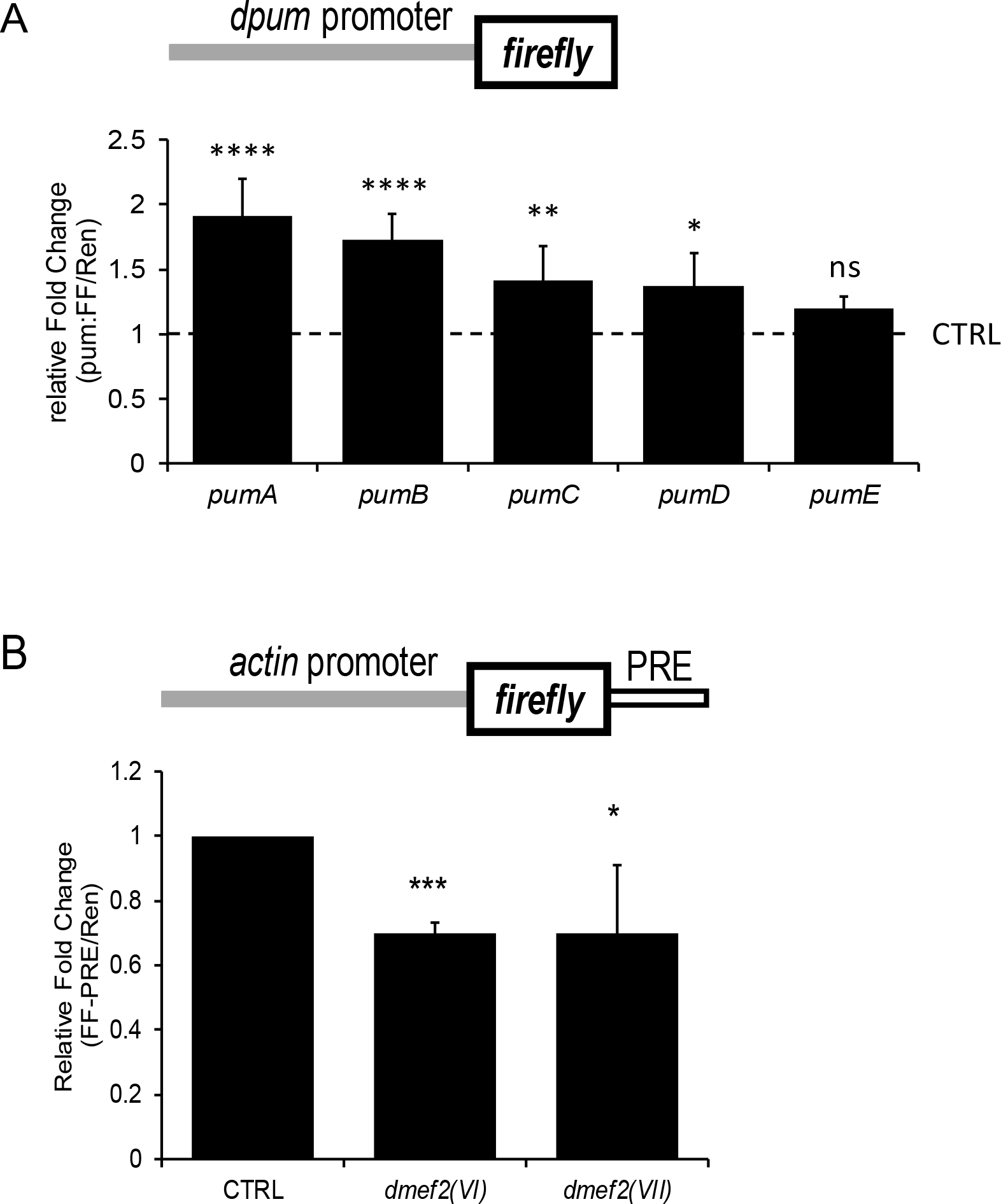
dMef2 binding elements are required for *dpum* promoter transactivation and increased dPum activity. A *dpum* promoterfirefy-luciferase *(pum:FF)* constructs, containing different numbers of dMef2 binding motifs *(pumA-E*, 4, 3, 3, 2 and 0, respectively), were cotransfected with *dmef2(VII)* and a renilla-luciferase gene *(Ren*, loading control, driven by *actin* promoter) in S2R+ cells. Expression of *dmef2* resulted in 1.9 ± 0.3, 1.7 ± 0.1, 1.4 ± 0.3, 1.4 ± 0.3 and 1.2 ± 0.1-fold increase in promoter activity *(pumA-E*, respectively) compared to control (set at 1: each construct expressed in the absence of *dmef2(VII))* (n = 5 independent transfections). B dPum protein activity can be measured using an *actin* promoter driven*firefly-* luciferase (FF) reporter gene containing Pumilio Response Elements (PRE) in the 3’ UTR (FF-PRE). Increased dPum is sufficient, through binding the PREs and inhibiting translation, to reduce FF activity. An identical renilla-luciferase (Ren) reporter, but lacking the PRE sites (and thus not affected by dPum), is co-expressed to allow ratiometric determination of activity. Expression of *dmef2(VI)* or *dmef2(VII)* isoforms in S2R+ cells, that express both reporters, significantly reduces the ratio of FF-PRE/Ren to 0.7 ± 0.04 and 0.7 ± 0.21, respectively (control (CTRL), no *dmef2* expression, set as 1) (n = 5 independent transfections). Data information: Data are presented as mean ± s.d. *P < 0.05, **P < 0.01, ***P ≤ 0.001 and ****P ≤ 0.0001. ns: not significant. (ANOVA with Bonferroni’s post-hoc).

### dMef2 regulates Pumilio expression level

To confirm our observations of transcriptional regulation of *dpum* by dMef2, we determined effect to dPum protein level. Because of the poor performance of commercially-available anti-Pum antibodies for Western Blotting in *Drosophila* (our unpublished data), we previously developed a monitor of dPum protein activity (Lin et al., 2017). Essentially, we constructed an *actin* promoter driven firefly-luciferase reporter gene (FF-PRE), containing PREs, (Pumilio Response Elements) in the 3’ UTR. Increased dPum is sufficient, through binding the PREs and inhibiting translation, to reduce FF activity. An identical renilla-luciferase *(Ren)* reporter, but lacking the PRE sites (and thus not affected by dPum), is co-expressed to allow ratiometric determination of activity (to compensate for batch differences between construct expression). Overexpression of *dmef2(VI)* or *dmef2(VII)* isoforms in S2R+ cells, that express both reporters, significantly reduces the ratio of FF-PRE/Ren to 0.7 ± 0.04 and 0.7 ± 0.21, respectively, (control, no *dmef2* expression, set as 1, *p* = 0.00013 and 0.03, respectively, Fig. 4B). We conclude that increasing *dmef2* is sufficient to increase dPum protein, in addition to transcript abundance.

### p300 supresses dMef2-mediated *pumilio* transactivation

Our prior work has shown levels of dPum and Pum2 (in fly and rat, respectively) are sensitive to neuronal activity: increasing as levels of synaptic excitation increase (Driscoll et al., 2013, Mee et al., 2004). It was expected, therefore, that *dmef2* would show activity-dependent transcription. Similar to mammals, ingestion of the proconvulsant picrotoxin (PTX) by larvae is sufficient to increase synaptic excitation and induce a seizure-like state (Stilwell et al., 2006). Therefore, we performed RT-qPCR to examine *dmef2* transcript expression in CNS taken from PTX-fed larvae. We did not, however, observe a significant fold-change (0.97 ± 0.06, n = 5, *p* > 0.05) compared to vehicle control (set at 1). This lack of effect is indicative that expression of *dmef2*, in *Drosophila*, is not activity-dependent. Moreover, if dMef2 regulates transcription of *dpum*, this would likely be through additional, and activity-dependent, regulators.

p300 is a reported co-regulator of Mef2A in mammal (De Luca et al., 2003) and we identify potential binding sites (GGGAG) for this protein in the *dpum* promoter (Chen & Hung, 1997, Rikitake & Moran, 1992), (see Fig. 1). A previous RNA-seq analysis, between isolated CNS taken from wildtype (WT) larvae and WT larvae fed PTX, identified *Drosophila p300* to be significantly down-regulated by enhanced synaptic excitation (214 ± 10 vs 163 ± 2 counts per million, WT vs. WT fed PTX, n = 3, *p* = 0.001) (Lin et al., 2017). To confirm this observation, we performed RT-qPCR to examine *p300* transcript abundance in CNS taken from PTX-fed WT larvae: *p300* transcript expression is downregulated to 0.87 ± 0.04 (n = 5, *p* = 0.0026) compared to vehicle control (set at 1).

To test how p300 contributes to transcriptional regulation of *dpum* expression, we coexpressed *p300* and *pumA:FF* in S2R+ cells. Overexpression of *p300* was sufficient to reduce *pumA* promoter activity to 0.76 ± 0.07, *(pumA:FF* alone set at 1, n = 5,*p* = 0.005). Moreover, dMef2-mediated activation of *pumA:FF* is abolished when coexpressed with *p300* (0.75 ± 0.04, n = 5, *p* = 0.0034) (Fig. 5A). Co-transfection with varied doses of *p300* showed a clear dose-dependent suppression of dMef2-mediated transactivation (2.79 ± 0.22, 1.8 ± 0.17, 1.53 ± 0.03, 1.19 ± 0.05 and 0.76 ± 0.07, *p300* plasmid: 0, 10, 20, 50 and 100 ng, respectively, *p* = 4.2×10^-15^) (Fig. 5B). Pretreatment of S2R+ cells with *p300* dsRNA enhanced *pumA:FF* promoter activity to 2.12 ± 0.22 (n = 5, *p* = 0.0001) (Fig. 5A). Finally, overexpression of *dmef2*, in the presence of *p300* dsRNA, further increased *pumA:FF* activity to 4.41 ± 0.74 (n = 5, *p* = 0.0001) (Fig. 5A). Collectively, these data suggest that p300 negatively regulates the *dpum* promoter. However, it does not discriminate whether p300 acts through direct binding to the *dpum* promoter.

**Figure 5.**
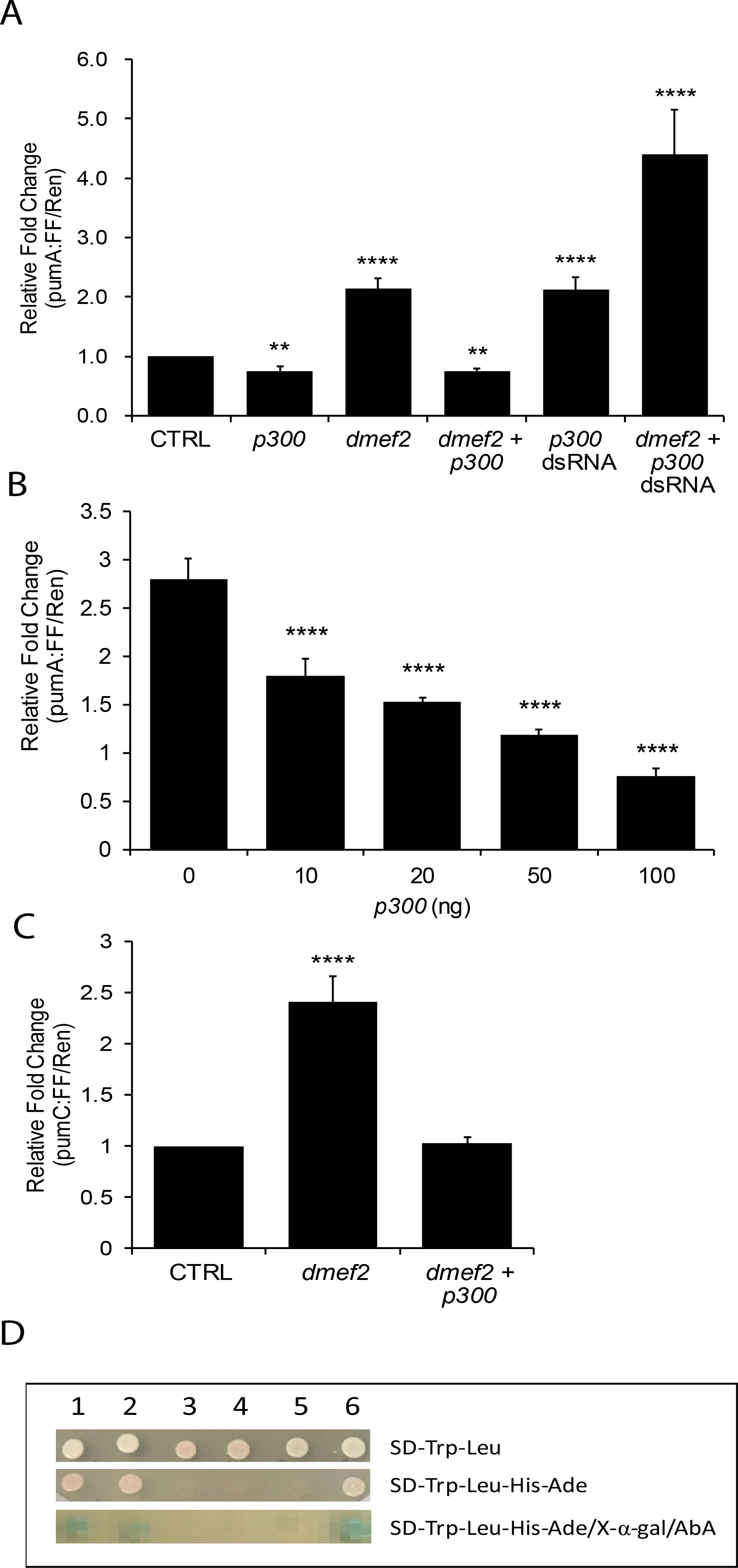
p300 represses dMef2-mediated transactivation of the *dpum* promoter. A The *pumA* promoterfirefy-luciferase *(pumA.FF)* reporter was co-transfected with *p300* and a renilla-luciferase gene *(Ren*, loading control, driven by *actin* promoter) in S2R+ cells. Expression of *p300* reduced *pumA* promoter activity to 0.76 ± 0.07, (control (CTRL), pumA:FF/Ren alone set at 1). By comparison, expression of *dmef2(VII)* resulted in a 2.14 ± 0.18-fold increase in luciferase activity compared to control. The activity of *dmef2(VII)* is abolished when co-expressed with*p300* (0.75 ± 0.04). Pre-treatment of S2R+ cells with *p300* dsRNA enhanced *pumA* promoter activity to 2.12 ± 0.22. Overexpression of *dmef2*, in the presence of *p300* dsRNA, further increased *pumA* promoter activity to 4.41 ± 0.74 (n = 5 independent transfections). B Co-transfection of *pumA:FF* with increasing doses of *p300* showed a clear dose- dependent suppression of dMef2(VII)-mediated transactivation (2.79 ± 0.22, 1.8 ± 0.17, 1.53 ± 0.03, 1.19 ± 0.05 and 0.76 ± 0.07, *p300* plasmid: 0, 10, 20, 50 and 100 ng, respectively, pumA:FF/Ren alone set at 1) (n = 5 independent transfections). C Co-transfection of *pumC:FF* (which lacks consensus p300 binding sequences) with *dmef2(VU)* resulted in a 2.41 ± 0.2-fold increase in luciferase activity compared to control (CTRL, pumC:FF/Ren alone set at 1). This enhancement is abolished when co-expressed with *p300* (0.75 ± 0.04) (n = 5 independent transfections). D Yeast two-hybrid assay shows a protein-protein interaction between dMef2 and p300. The bait plasmid pGBKT7-p300 (BD-p300) and the prey pGADT7-dmef2 (AD- *dmef2(VI)* or *AD-dmef2(VII))* were co-transformed into a Y2HGold yeast strain. Coexpression of 1. BD-p300/AD-dMef2(VI) or 2. BD-p300/AD-dMef2(VII) resulted in activation of all four GAL4-responsive markers, *HIS3, ADE2, AUR1-C* and *MEL1* reporters in Y2HGold. The negative control groups, 3. BD-lam/AD-dMef2(VI), 4. BD-lam/AD-dMef2(VII) and 5. BD-p300/AD-T, were not able to activate reporter gene expression in Y2HGold. 6. BD-p53/AD-T was used as a positive control. AbA: Aureobasidin A, BD: GAL4 DNA binding domain, AD: GAL4 activation domain, SD: synthetic dropout selective media, Lam: Lamin C, T: large T antigen. Data information: in (A-C), data are presented as mean ± s.d. **P ≤ 0.01 and ****P ≤ 0.0001. (ANOVA with Bonferroni’s post-hoc).

To test whether binding of p300 to the *dpum* promoter is required for repression of dMef2-dependent *dpum* transactivation, we expressed *p300* and *dmef2* with *pumC:FF*, which lacks consensus p300 binding sequences (see Fig. 1). Whilst expression of dMef2 enhanced *pumC:FF* activity (2.41 ± 0.2, *pumC:FF* alone set at 1, n = 5, *p* = 1.4×10^-6^), co-expression of *p300*, however, was still sufficient to significantly reduce dMef2-mediated transactivation (1.0 ± 0.1, n = 5, *p* > 0.05, Fig. 5C). Thus, direct binding of p300 to the *dpum* promoter is seemingly not required for repression of dMef2-mediated *dpum* transactivation, indicative of a protein-protein interaction between p300 and dMef2. To demonstrate a physical interaction between p300 and dMef2, we performed a yeast two-hybrid assay. The bait plasmid pGBKT7-p300 (BD-p300) and the prey *pGADT7-dmef2 (AD-dmef2(VI)* or *AD-dmef2(VII))* were cotransformed into a Y2HGold yeast strain. Co-expression of BD-p300 with AD- dMef2(VI) or AD-dMef2(VII) resulted in activation of all four GAL4-responsive markers, *HIS3, ADE2, AUR1-C* and *MEL1* reporters in Y2HGold. Conversely, coexpression of control groups, BD-p300/AD-T (large T antigen) or AD-dMef2/BD-Lam (Lamin C), was not able to activate reporter gene expression, indicative that the auto-activation of BD-p300 and AD-dMef2 were negligible (Fig. 5D). This result strongly suggests that p300 directly binds dMef2.

### p300 represses *pumilio* promoter activity *in vivo*

To test *dpum* promoter activity *in vivo*, we generated a transgenic *pumC-GAL4* fly and mated it with *UAS-luciferase* (UAS-luc) (Markstein et al., 2008). The resultant 3^rd^ instar larvae, expressing *pumC-GAL4>luc* showed a significant increase (2.1 ± 0.2- fold) in luc activity compared to control (UAS-luc line, set at 1, *p* = 1.3×10^-5^, *t*-test, n = 10). To test how the *pumC* promoter responds to increased synaptic excitation, we established a stable line of *pumC-GAL4>luc* and raised larvae on food containing PTX (1 µg/ml). This resulted in a 1.7 ± 0.4-fold increase in luc compared to vehicle control (i.e. no PTX, set at 1, *p* = 0.003, n = 10, *t*-test). This result not only confirms our previous observations that *dpum* expression is regulated by membrane depolarization (Mee et al., 2004), but transfers our capability to measure *dpum* promoter activity to *in vivo*.

Overexpression of *dmef2*, by crossing *pumC-GAL4>luc* with UAS-dmef2, resulted in a 2.0 ± 0.3-fold increase in luc expression compared to *pumC-GAL4>luc* crossed with control UAS-GFP (set at 1, *p* = 0.0005, n = 10, Fig. 6). This confirms that the *pumC* promoter is transactivated by dMef2 *in vivo*. To validate our observation that p300 represses *dpum* expression, we crossed *pumC-GAL4>luc* with *UAS-p300* or UAS- p300^RNAi^. Overexpression of *p300* resulted in a 0.4 ± 0.2-fold reduction in luc activity (*p* = 0.0005, n = 10), whilst p300-knockdown increased luc expression by 1.3 ± 0.4- fold (*p* = 0.02, n = 10) (Fig. 6). p300 contains histone acetyltransferase (HAT) activity (Ogryzko et al., 1996). To test if HAT activity is required for the repression of *dpum* transactivation, we crossed *pumC-GAL4>luc* with UAS-p300 ^*F2161A*^, a mutant with an abolished HAT activity (Ludlam et al., 2002). Overexpression of *p300^F2161A^* still resulted in a 0.3 ± 0.1-fold reduction in luc activity (*p* = 0.0003, n = 10) (Fig. 6), indicating that p300-dependent repression of *dpum* expression is independent of HAT activity. Thus, we validate *in vivo* that dMef2 is sufficient to transactivate the *dpum* promoter and that this activity is negatively regulated by p300.

**Figure 6.**
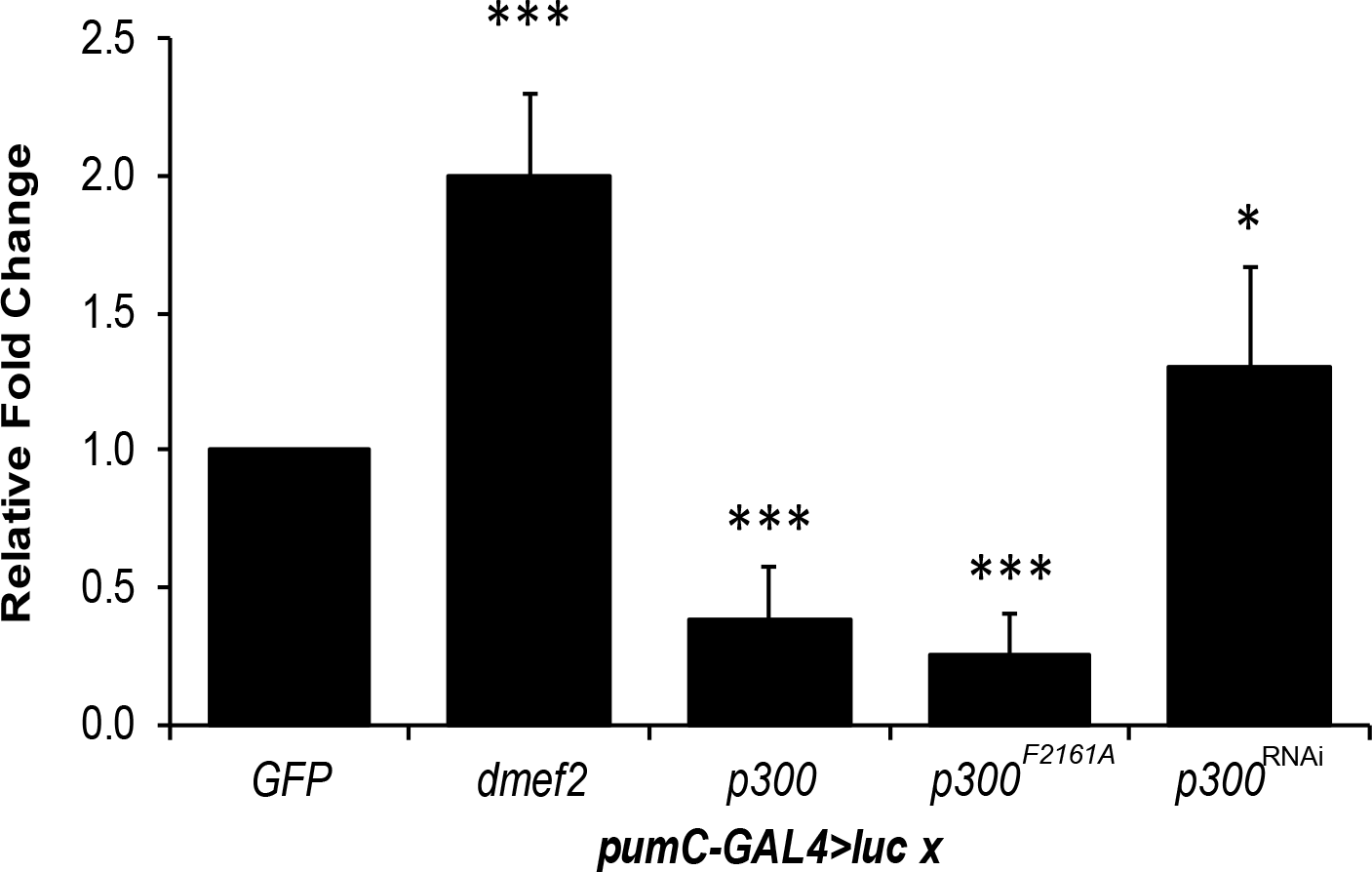
dMef2 and p300 regulate *dpum* promoter activity *in vivo*. Expression of *dmef2, p300* or *p300^F2161A^*, respectively, resulted in a 2.0 ± 0.3, 0.4 ± 0.2 and 0.3 ± 0.1-fold change in luciferase (luc) activity compared to control *(pumC- GAL4>luc* cross with UAS-GFP *(GFP)*, set at 1). Knock-down of *p300* expression, achieved by crossing UAS-p300^RNAi^ *(p300^RNAi^)*, resulted in a 1.3 ± 0.4-fold increase in luc activity (n = 10 independent sample collections). Data information: data are presented as mean ± s.d. *P ≤ 0.05 and ***P ≤ 0.001. (ANOVA with Bonferroni’s post-hoc).

## Discussion

Pum is a key regulator of neuronal firing-rate homeostasis that maintains action potential firing within physiologically-appropriate limits (Giachello & Baines, 2017, Mee et al., 2004, Muraro et al., 2008). Despite this critical role, little is known concerning how Pum levels are regulated. Its involvement with homeostasis suggests that *pum* transcription will be governed by an activity-dependent process (Mee et al., 2004). In this study, we characterised the *dpum* promoter region and identified both Mef2 and p300 to be part of an activity-dependent regulatory mechanism. The complete regulatory mechanism is, however, likely to be more complicated based on our identification of binding motifs for 114 putative transcription factors within a 2-kb region upstream of the *dpum* transcription start site. A series of deletion constructs show that *pumA, pumB* and *pumC* promoters (compose of 2000, 1434 and 578 nucleotides, respectively) exhibit promotor activity, whilst *pumD* and *pumE* (consist of 312 and 189 nucleotides, respectively) result in a reduced, but still significant, activity. The *pumE* promoter, the shortest construct we tested, contains binding motifs for 15 transcription factors (e.g., Pbx-1, BR-C Z4, Zen and Lhx3a) and is, as we show, still capable of driving *FF* expression in S2R+ cells. Further truncations will be required to identify the minimal promoter region for *dpum.*

Multiple dMef2 binding motifs are located within the 2-kb *dpum* promoter region and manipulating *dmef2* expression is sufficient to influence promoter activity. This strongly suggests that *dpum* expression is regulated, at least in part, by dMef2. *Dmef2,* which is encoded by a single gene, contains 15 exons. Of these exons, 4 are subject to alternative splicing and generate at least 12 splice variants (*dmef2(I-XI)* and *dmef2(mini))*. All splice isoforms transactivate the *dpum* promoter and, notably, exon 10 containing isoforms result in greater activation compared to variants lacking this exon. This differential activity may be indicative that the level of *dpum* transactivation can be fine-tuned through altering of dMef2 isoform expression. In muscle, however, it is the expression level of dMef2, rather than isoform expression, which is seemingly more important for cellular differentiation (Gunthorpe et al., 1999). This conclusion was reached, however, without consideration of exon 10 lacking isoforms, because splicing of exon *10* was not previously observed (Gunthorpe et al., 1999, Taylor et al., 1995). Our bioinformatics shows that Mef2 binding sites are also conserved in the mammalian (e.g. human and mouse) *pum2* promoter region. Interrogation using the Harmonizome search engine followed by ChIP-X enrichment analysis (ChEA) of transcription factor targets database, identifies 23 transcription factors, including p300 and Mef2A, associate with the *pum2* promoter region (Lachmann et al., 2010, Rouillard et al., 2016). A genome-wide tiling array (ChIP-Chip) analysis similarly identifies *dpum* as a target of dMef2 (Sivachenko et al., 2013). Our analysis of the promoter region (-2000 to +1, set transcription initiation at +1), in both mouse and human *pum2*, identifies 5 and 6 Mef2 elements, respectively (Table EV2 and EV3). Thus, it seems highly likely that Mef2 is a direct regulator of *pum* transcription in both insects and mammals. This extends the activity of this transcription factor in addition to its reported inhibitory control of *pum2* transcript abundance via up-regulation of miR-134 (Fiore et al., 2009, Fiore et al., 2014). It will be important to understand the relative efficacy of the different Mef2 splice variants for their ability to regulate expression level of miR-134.

A lack of effect on *dmef2* expression level following exposure to PTX is indicative that this factor does not form a primary link between neuron membrane depolarization and altered expression of *dpum.* Our studies instead spotlight p300. p300 contains HAT activity and is also an accessory protein that interacts with other transcription factors to function as either a co-activator or as a repressor. For example, in *Drosophila*, p300 interacts with Dorsal, Mad and Cubitus interruptus, to co-activate *Toll, decapentaplegic* and *Hedgehog* signalling pathways, respectively (Akimaru et al., 1997a, Akimaru et al., 1997b, Waltzer & Bienz, 1999). By contrast, T-cell factor (TCF)-mediated Wnt/Wingless signalling is repressed by p300 acetylation (Waltzer & Bienz, 1998). In mammals, p300 has been reported to bridge the complex of thyroid hormone receptor-retinoid X receptor-Mef2A to abrogate transactivation of α-myosin heavy chain gene promoter activity when an inhibitor, adenovirus E1A for example, is recruited (De Luca et al., 2003). Here we show p300 acts as a repressor of dMef2- mediated transactivation of *dpum.* We further show that this effect is likely achieved through direct binding of p300 to dMef2. This result mirrors the reported physical interaction between the cysteine/histidine-rich region 3 (CH3) domain of p300 and Mef2A/Mef2C (through MASD/Mef2 domains) in mammals (De Luca et al., 2003, Sartorelli et al., 1997).

Our results are consistent with increasing neuronal depolarization negatively regulating expression of p300 which, in turn, allows increased dMef2-mediated transactivation of *dpum.* Expression of *p300* is known to be activity-dependent and is similarly downregulated in pilocarpine (increased acetylcholine signalling) treated mouse hippocampus (Hansen et al., 2014). Expression of *mef2* has also been reported to be regulated by increased synaptic excitation in mammals (Mao et al., 1999), an observation we could not validate in *Drosophila.* We cannot rule out, however, that additional post-transcriptional and/or post-translational modifications, including alternative splicing and/or phosphorylation of *dmef2* might influence its activity. In this regard, a report of Mef2 activation by Ca^2+^-activated dephosphorylation, via calcineurin, is particularly attractive (Flavell et al., 2006). This is because Ca^2+^ entry across the neuronal membrane is widely regarded as an initial reporter of neuronal activity in homeostatic mechanisms (Cudmore & Turrigiano, 2004, Gunay & Prinz, 2010, O’Leary et al., 2010). Alternative post-transcriptional modifications of Mef2 activity have been reported. For example, p38 mitogen-activated protein kinase (p38-MAPK) induced phosphorylation of Mef2C is critical for activation of Mef2 target genes (Han et al., 1997, Mao et al., 1999).

Up-regulation of *dpum* transcript expression in late stage 17 *Drosophila* embryos, due to increased synaptic excitation, was previously reported (Mee et al., 2004). Intriguingly, the analysis of transcript expression in 3^rd^ instar CNS between wildtype and wildtype raised on food containing the PTX revealed a significant reduction of *dpum* transcript expression (Lin et al., 2017). These paradoxical results might be a result of *dpum* auto-regulation. The *dpum* transcript contains multiple PRE motifs in its 3’UTR region (Chen et al., 2008, Gerber et al., 2006). Similarly, human *PUM1* and *PUM2* are also potential targets of PUM protein (Bohn et al., 2018). In this study, we used a *dpum* promoter to drive *firefly-luc* expression, which lacks PRE motifs, and showed enhanced *dpum* promoter activity in 3^rd^ instar larvae raised on food containing PTX. This result validates that the *dpum* promoter is responsive to levels of synaptic activity and also provides additional evidence to suggest that Pum regulates its own expression through negative-feedback.

In summary, we show that regulation of *dpum* expression is mediated by an interaction between dMef2 and p300, the latter being an activity-dependent negative regulator. Under ‘normal’ synaptic excitation, p300 is more abundant and binds to dMef2 to inhibit transactivation of *dpum*. Increased synaptic excitation reduces *p300* expression which, in turn, releases dMef2 from inhibition. The increase in dPum protein translationally represses *paralytic* (*voltage-gated sodium channel*) mRNA to achieve a homeostatic reduction in action potential firing (the schematic mechanism is shown in Fig. 7). It follows that inhibition of p300 would be predicted to be anticonvulsant (mirroring increased dPum activity). Indeed, treatments (genetic or pharmacological) that elevate dPum activity are potently anticonvulsive in *Drosophila* seizure mutants (Lin et al., 2017). Moreover, seizures in flies, rodents and human are associated with a decrease in dPum or Pum2 activity, respectively (Follwaczny et al., 2017, Lin et al., 2017, Siemen et al., 2011, Wu et al., 2015). Thus, whilst drug interventions that directly active Pum may be difficult to achieve in the clinic, inhibition of p300 may represent a more achievable route to better control epilepsy.

**Figure 7.**
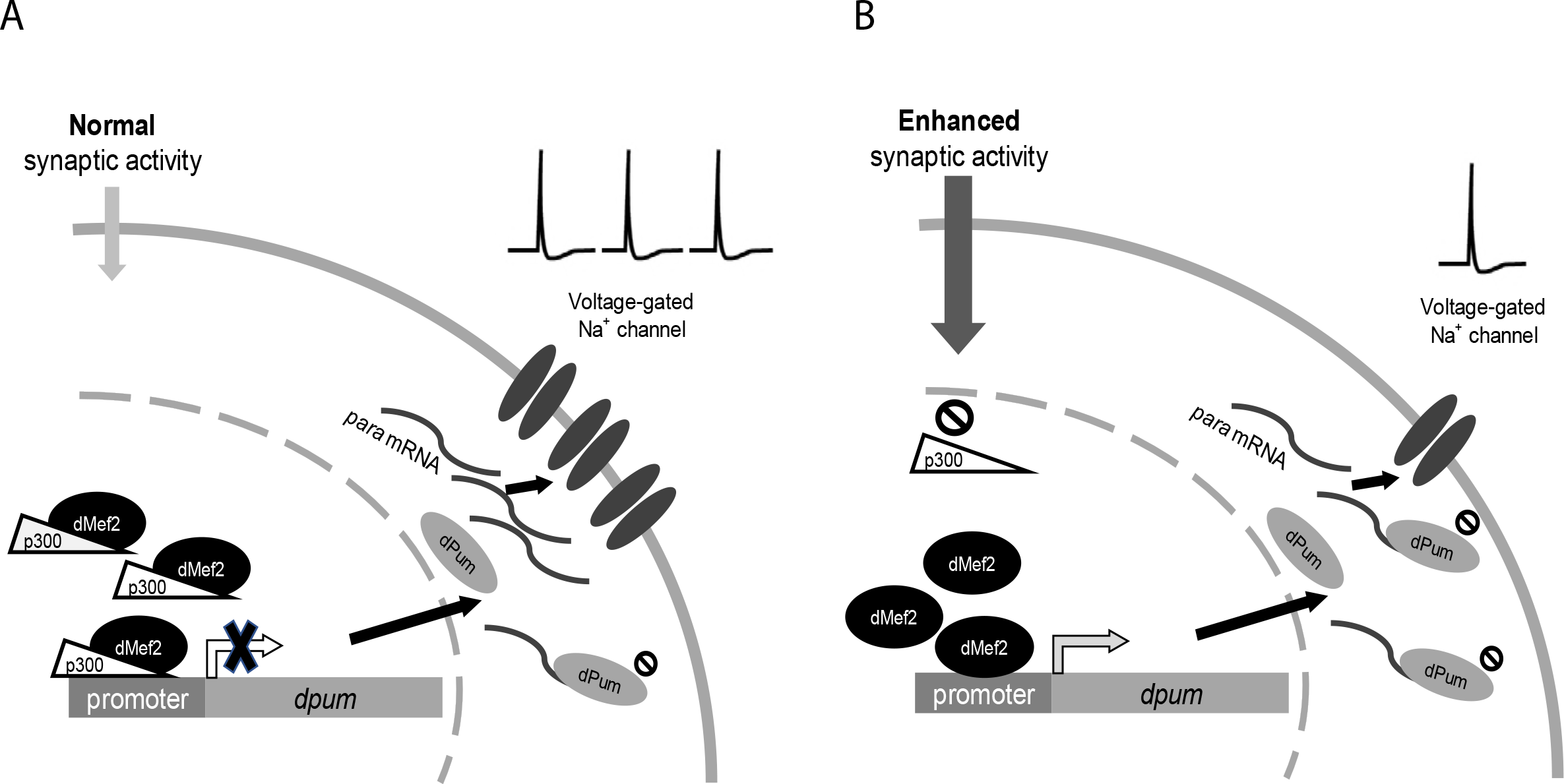
Transcription of *dpum* is co-regulated by dMef2 and p300. A Under ‘normal’ synaptic excitation, p300 is more abundant and binds to dMef2 to inhibit the transactivation of *dpum.* B Increasing exposure to synaptic excitation is sufficient to downregulate expression of *p300* resulting in the release of dMef2 from inhibition. This facilitates transactivation of *dpum.* Increased dPum protein can, in turn, translationally repress *paralytic* mRNA *(para*; *voltage-gated sodium channel)* to achieve a homeostatic reduction in action potential firing.

## Materials and Methods

### Cloning of expression plasmids

#### *dpum* promoter

*Pumilio (dpum*, CG9755) genomic sequence was obtained from FlyBase (http://flybase.org). Genomic DNA from Canton-S wildtype was extracted in 50 μl extraction buffer (10 mM Tris-HCl, 1 mM EDTA, 25 mM NaCl and 200 μg/ml proteinase K) with incubation at 37°C for 30 min. *Dpum* promoter constructs were amplified by PCR (Phusion High-Fidelity DNA Polymerase, New England Biolabs, Hitchin, UK) that consisted of the following in a total volume of 50 μl: 20 pmol primers, dNTPs at 0.2 mM, and 1X Phusion HF buffer with 1.5 mM Mg^2+^. The forward and reverse primers introduced a *Kpn* I and a *Xho* I sites at the 5’ and 3’ end of promoter, respectively. Cycling conditions were: initial denaturation at 98°C for 5 min; 35 cycles of 98°C for 10 sec, 55°C for 20 sec and 72°C for 2 min 30 sec; a final extension step at 72°C for 10 min. The PCR product was digested with *Kpn* I and *Xho* I and ligated into pGL4.23 vector (Promega). The forward and reverse primer sequences are as follows (5’ to 3’): pumA (-2000 to +1), AATA**GGTACC**CGATGGCTCCGGCGCTGA and pumR: TATT**CTCGAG**GAACATTTAGTGTGACCGCAGCT. A series of deletion constructs for the *dpum* promoter were PCR amplified using forward primers, pumB (-1434 to +1), AATA**GGTACC**GACCGTCGGCTGGATCCGT, pumC (-578 to +1), AATA**GGTACC**ACATAGCTCGGAAAACGATTTCAAC, pumD (-312 to +1), ATAT**GGTACC**ATGGTTGTATTGATTCTTTATAT and pumE (-189 to +1), ATAT**GGTACC**GGCAACTAGTTAAATGCATTATAG and the reverse primer, pumR.

#### Amplification of *dmef2* splice variants and *p300*

Total RNA was extracted from 3^rd^ instar CNS of Canton-S. cDNA synthesis was carried out in a total volume of 20 μl using the manufacturer’s protocol (RevertAid First-Strand cDNA Synthesis kit; Thermo Fisher Scientific). *Dmef2* PCR was performed by using forward and reverse primers, which introduced a *Kpn* I and a *Xho* I site, respectively, and ligated to pAc5.1 expression vector (Thermo Fisher Scientific). Fifty-six plasmids from independent *E*. *coli* colonies were isolated and sequenced to identify splice variants of *dmef2*. The forward and reverse primer sequences are as follows (5’ to 3’): ATTA**GGTACC**GGAT AGGAAATCTGTTGCCATGG and ATTA**CTCGAG**CAGCTCGTGCCGGCTATGT. *p300 (nejire*, CG15319) was PCR-amplified with the primer pairs which introduced *Kpn* I and *Xba* I sites in the 5’ and 3’ end of the open reading frame, respectively. PCR product was ligated to pAc5.1 expression vector. The forward and reverse primer sequences are as follows (5’ to 3’): AATA**GGTACC**ATGATGGCCGATCACTTAGACG and AATA**TCTAGA**CTAGAGTCGCTCCACAAACTTG. All clones were checked by sequencing prior to expression analysis.

### Identification of transcription factor binding sites

Mouse and human *pum2* promoter sequences (-2000 to +1) were obtained from the National Centre for Biotechnology Information (NCBI:https://www.ncbi.nlm.nih.gov), mouse: GRCm38:12: 8672314:8674133 and Human: NC_000002.12:c20354428-20352429. Transcriptional elements and factors were predicted using the TRANSFAC models of MAPPER search engine (http://genome.ufl.edu/mapper/). Mammalian transcription factors associated with human and mouse *pum2* promoters were identified by the Harmonizome search engine (http://amp.pharm.mssm.edu/Harmonizome/).

### Luciferase assay

S2R+ cells (10^6^ cells in 100 μl of Schneider’s *Drosophila* Medium, Gibco) were treated with dsRNA (1 μg) in a 96-well plate (Corning^®^ Costar^®^) for 3 h, followed by co-transfection (Effectene, QIAGEN) of *dpum promoter-firefly* construct and *renilla* luciferase reporter (100 ng each) for a further 24 h. The transfection procedure is as described in the manufacturer’s instructions (QIAGEN). The luciferase assay was performed using the Dual-Luciferase reporter assay system (Promega). Briefly, 30 μl of transfected S2R+ cells were transferred to a well of a 96-well white plate (FluoroNunc^™^) and lysed with 30 μl of passive lysis buffer and then 30 μl Luciferase Assay Reagent II was added to measure firefly luciferase activity. This was followed by 30 μl of Stop & Glo^®^ to measure renilla luciferase activity. A GENios plate reader (TECAN) was used to measure luminescence. At least five independent transfections of each experiment were performed. Double-stranded RNA, *dmef2* (BKN27383) and *p300* (BKN21411), were obtained from the Sheffield RNAi Screening Facility (Sheffield, UK).

Luciferase activities of 3^rd^ instar larvae were measured using the Promega Steady-Glo Luciferase Assay Kit. Briefly, the *dpum* promoter-GAL4 line (see below for details) was crossed to attP24 *UAS-luciferase* flies (Markstein et al., 2008). Flies carrying the *UAS-luciferase* transgene alone were used for background controls. Three larvae were collected in 200 μl Promega Glo Lysis buffer for each sample, and 10 independent samples collected for each genotype. Larvae were homogenized, incubated at room temperature for 10 min, centrifuged for 5 min, and supernatant was transferred to a new tube. For luciferase assays, 30 μl of each sample was transferred to a well of a white-walled 96-well plate at room temperature, 30 μl Promega Luciferase reagent was added to each well and plates were incubated in the dark for 10 min. Luminescence was measured with a GENios plate reader (TECAN). The obtained values were normalized to total protein concentration, measured using the Bradford protein assay (Bio-rad).

### Quantitative RT-PCR

Quantitative RT-PCR was performed using a SYBR Green I real-time PCR method (Roche, LightCycler^®^ 480 SYBR Green I Master, Mannheim, Germany) as described in Lin et al. (2015). RNA was extracted from 20 3^rd^ instar CNSs using the RNeasy micro kit (QIAGEN, Hilden, Germany). PCR primers were designed with the aid of LightCycler Probe Design Software 2.0 (v1.0) (Roche). Primer sequences (5’ to 3’) used were: *actin-5C* (CG4027), CTTCTACAATGAGCTGCGT and GAGAGCACAGCCTGGAT; *dmef2*, TTCAAATATCACGCATCACCG and GCTGGCGTACTGGTACA; *p300*, GTTCTGGACTTCCCACG and TACTGGCTCATTTGCATGTAAC (forward and reverse, respectively). Relative gene expression was calculated as the 2^-ΔCt^, where ΔCt was determined by subtracting the average *actin-5C* Ct value from that of *dmef2* or *p300.*

### Fly stocks

Transgenic flies were generated using the PhiC31 integrase-mediated transgenesis system. A pumC-GAL4-hsp70 fragment was constructed in the pattB vector and microinjected (BestGene Inc. Chino Hills, CA, USA) into an attP-containing fly stock (Bloomington *Drosophila* Stock Centre stock no. 9748: y^1^ w^1118^; PBac{y^+^-attP-3B}VK00031). UAS-p300 (stock no. 32573) and *UAS-p300^F2161A^* (stock no. 32574) were obtained from Bloomington and UAS-p300^RNAi^ (stock no. 102885) was obtained from the Vienna *Drosophila* Resource Centre. The UAS-dmef2 was a gift from Dr. Michael Taylor (Cardiff University, Cardiff, UK). The attP24 *UAS-luciferase* stock was a gift from Dr. Norbert Perrimon (Howard Hughes Medical Institute, Boston, USA).

### Yeast two-hybrid assay

Both *p300* (bait) and *dmef2* (prey) were cloned to pGBKT7 and pGADT7 vector, respectively. All yeast strains and plasmids (pGBKT7, pGADT7, pGBKT7-53, pGADT7-T and pGBKT7-Lam) were obtained from Clontech as components of the MATCHMAKER two-hybrid system 2. The *Escherichia coli (E. coli)* strain *DH5α* was used to clone every shuttle plasmid. Auxotrophic selection plates are Synthetic Dropout (SD) medium supplemented with 0.67% yeast nitrogen base, 0.06% appropriate dropout amino acid mixture and 2% bacto-agar. The purified bait and prey plasmids were co-transformed into the Y2HGold strain using a lithium-acetate method according to the manufacturer’s instructions (Clontech) and were then cultured on SD/-Trp/-Leu agar plates. Approximately 2 mm of Y2HGold transformants were transferred to SD/-Trp/-Leu/-His/-Ade and SD/-Trp/-Leu/-His/-Ade/X-α-gal/Aureobasidin A plates at 30 °C for 2∼3 d. The host *Saccharomyces cerevisiae* strain Y2HGold genotype is as follows: *MATa, trp1-901, leu2-3, 112, ura3-52, his3-200, gal4Δ, gal80Δ, LYS2:: GAL1_UAS_-Gal1_TATA_-His3, GAL2_UAS_-Gal2_TATA_-Ade2 URA3::MEL1_UAS_-MEL1_TATA_ AUR1-CMEL1.* In mammals, proteinprotein interaction between the cysteine–histidine-rich region 3 (CH3) domains containing C-terminus of p300 and Mef2A/Mef2C (through MASD/Mef2 domains) has been reported (De Luca et al., 2003, Sartorelli et al., 1997). Therefore, p300_301-3276_ (2976 amino acids, from 301^th^ to the end of C-terminus (3276^th^)) and full length dMef2 were cloned. p300 bait, BD-p300 DNA fragment was released from p300/pAc5.1 using *Nde* I and *Xba* I (filling the sticky end to blunt end with Klenow) and ligating into pGBKT7 *(Nde* I and *Sma* I). dMef2 prey, *AD-dmef2* DNA fragment was released from *dmef2(VI)/pAc5.1* or *dmef2(VII)/pAc5.1* using *Kpn* I (filling the sticky end to blunt end with Klenow) and *Xho* I and ligating into pGADT7 (*BamH* I (filling the sticky end to blunt end with Klenow) and *Xho* I).

### Statistics

Statistical significance between group means was assessed using either a Student’s *t*-test (where a single experimental group is compared to a single control group) or a one-way ANOVA followed by Bonferroni’s post-hoc test (multiple experimental groups). Data shown is mean ± standard deviation (s.d.)

## Acknowledgements

The authors thank Miaomiao He and Yuen Ngan Fan for excellent technical support and Dr. Carlo Giachello for help with production of figure 7. We thank Norbert Perrimon for the gift of *UAS-luciferase* flies. This work was supported by funding to RAB from the Biotechnology and Biological Sciences Research Council (BB/L027690/1). Work on this project benefited from the Manchester Fly Facility, established through funds from the University and the Wellcome Trust (087742/Z/08/Z).

## Author contributions

W.-H.L. and R.A.B. designed research; W.-H.L. performed research and analysed data; W.-H.L., and R.A.B. wrote the paper.

## Conflict of interest

No competing interests declared.

## Expanded View Figure legends

**Figure EV1.**
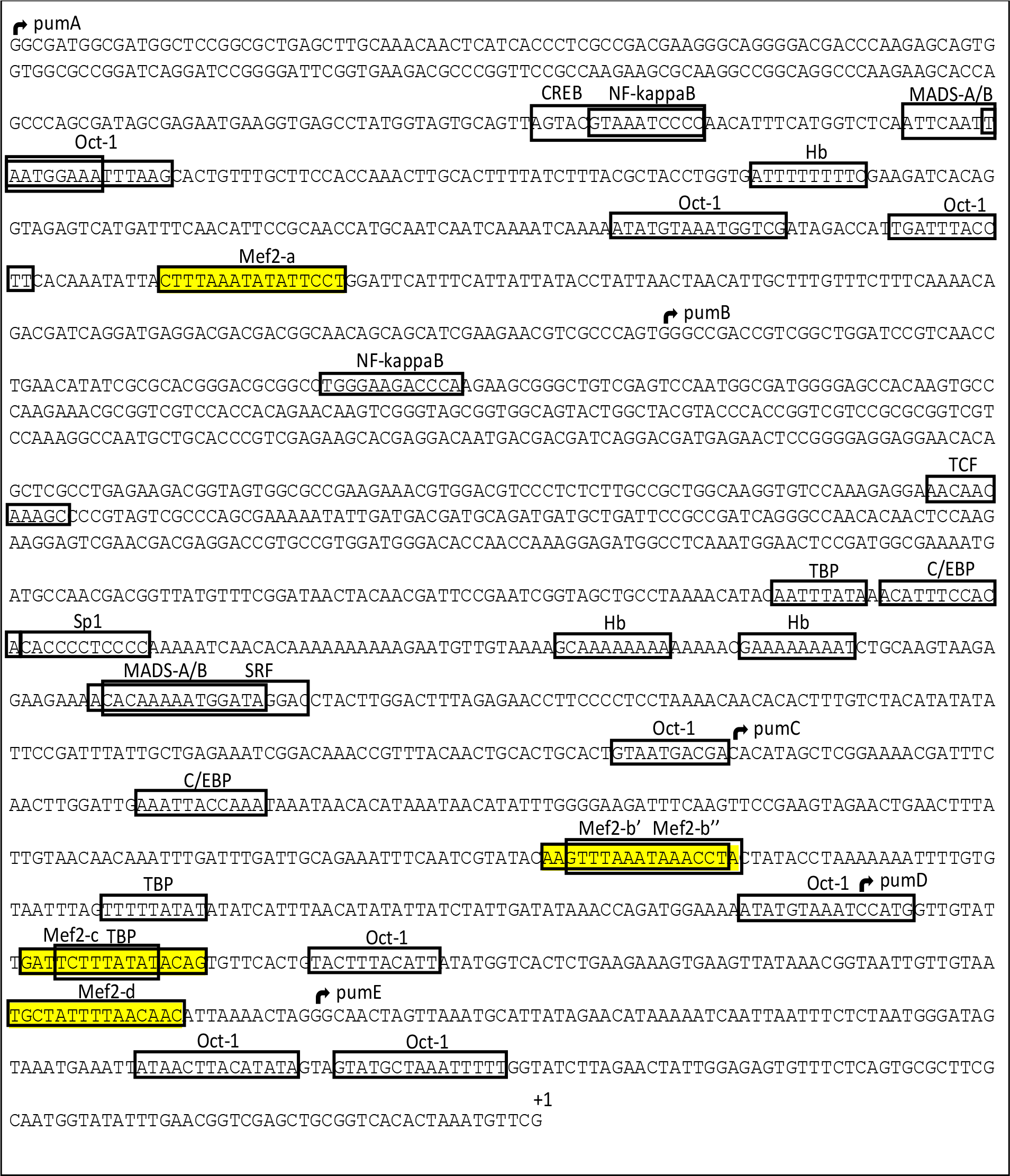
Putative transcription factors identified in the *dpum* promoter. A 2-kb region upstream of the transcription start site of *dpum* was interrogated using transcription factor databases (e.g. TRANSFAC model, MAPPER) (Marinescu et al., 2005). In total 114 putative transcription factors within the region between -2000 and +1 (transcription initiation marked as +1) were identified. Putative transcriptional elements including, Sp1, TBP, C/EBP, Oct-1, Mef2, MADS-A/B, Hb, NF-kappaB, TCF, CREB and SRF are highlighted. Four Mef2 binding sites, Mef2-a, -b, -c and -d, are highlighted in yellow. Two predicted Mef2 binding motifs, Mef2-b’ and Mef2-b’’ overlap and, therefore, we count this as one site (Mef2-b). The initiation of each *dpum* promoter fragment, *pumA-E*, are indicated as black arrows.

**Table EV1.**
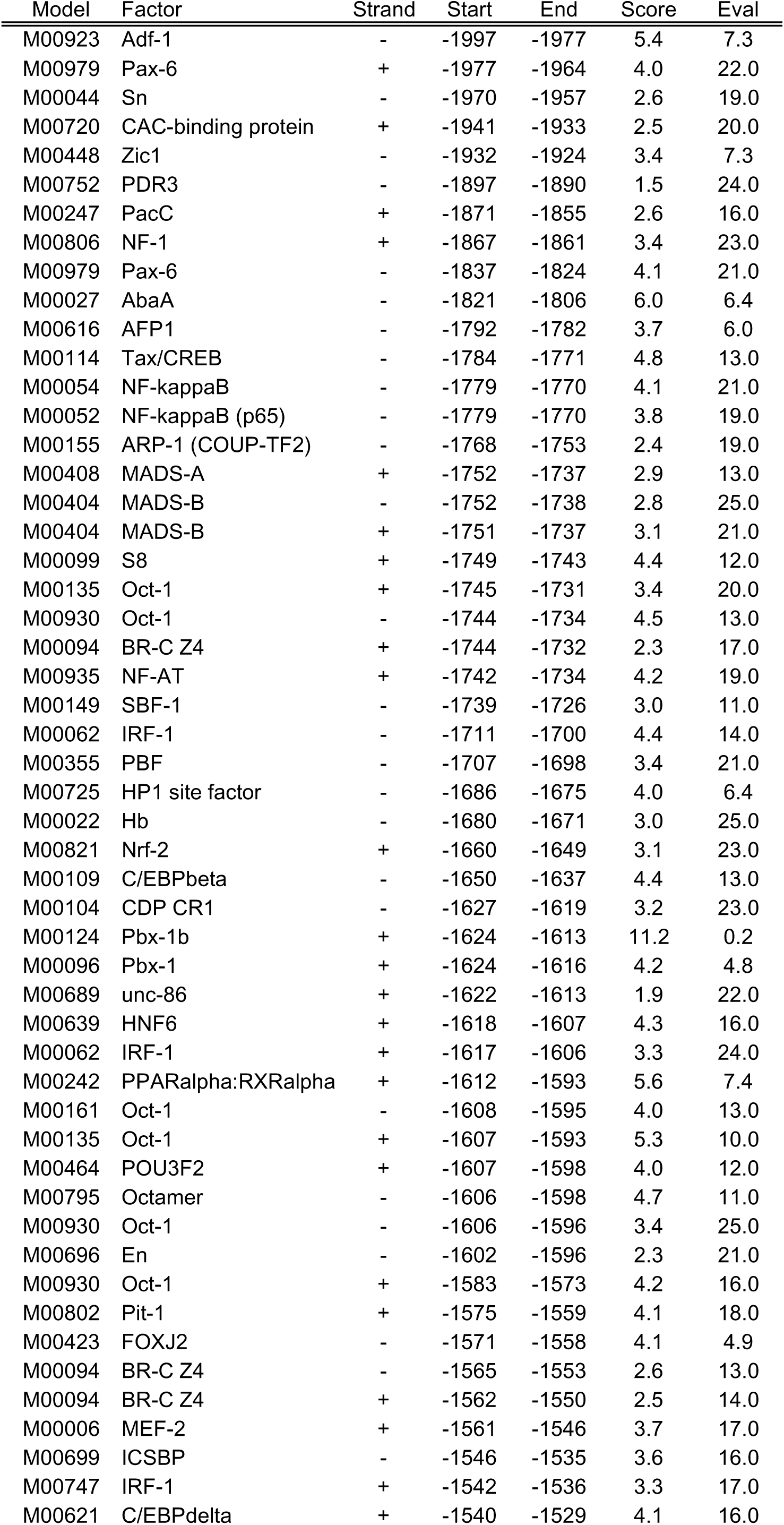
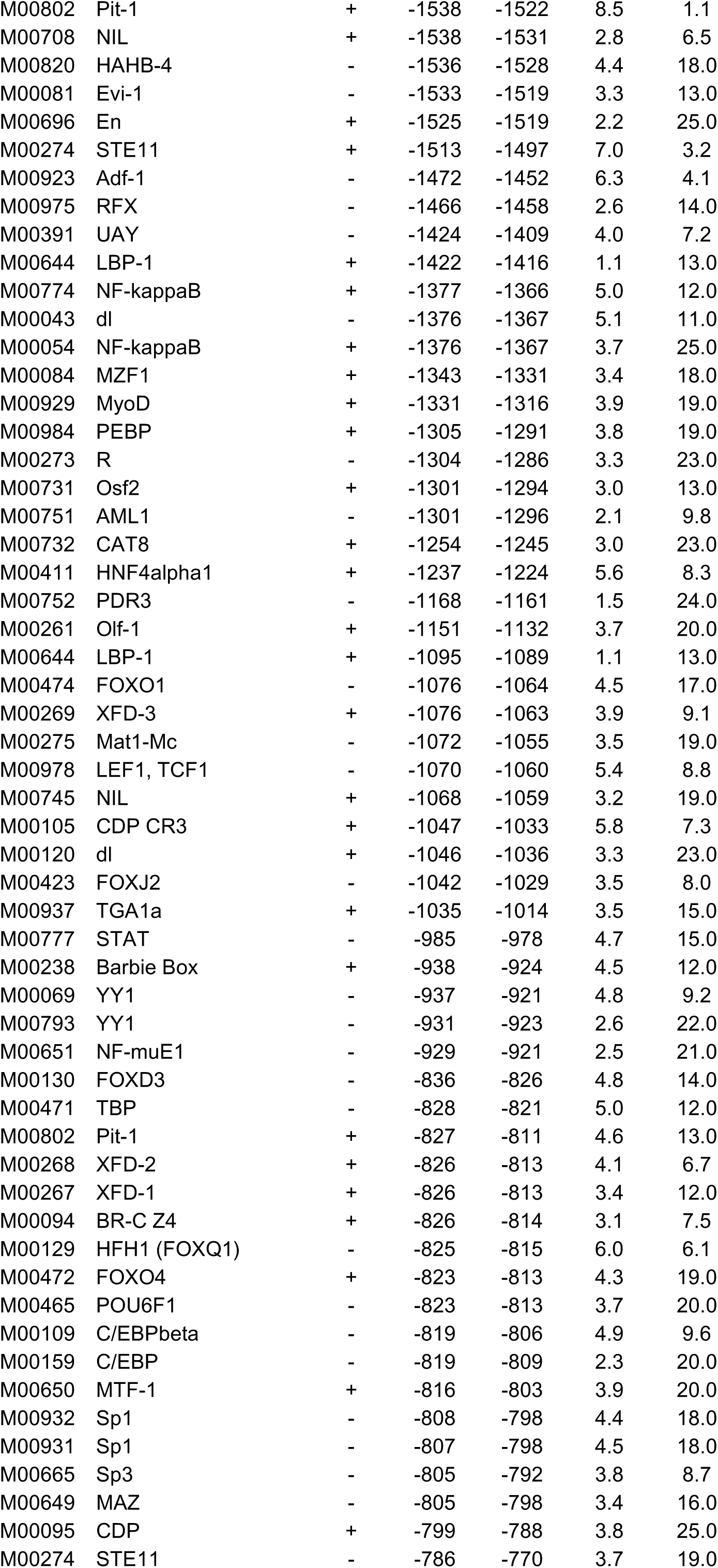
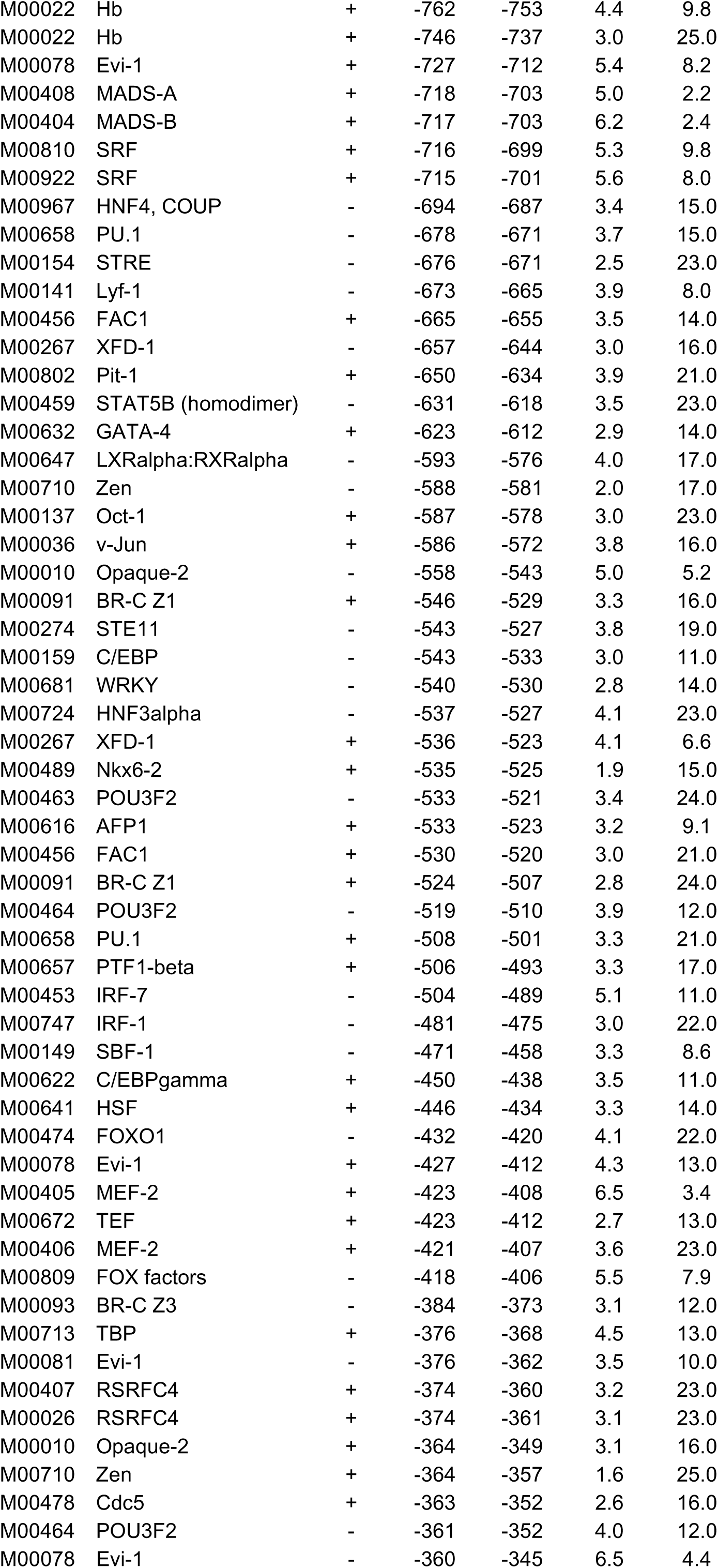
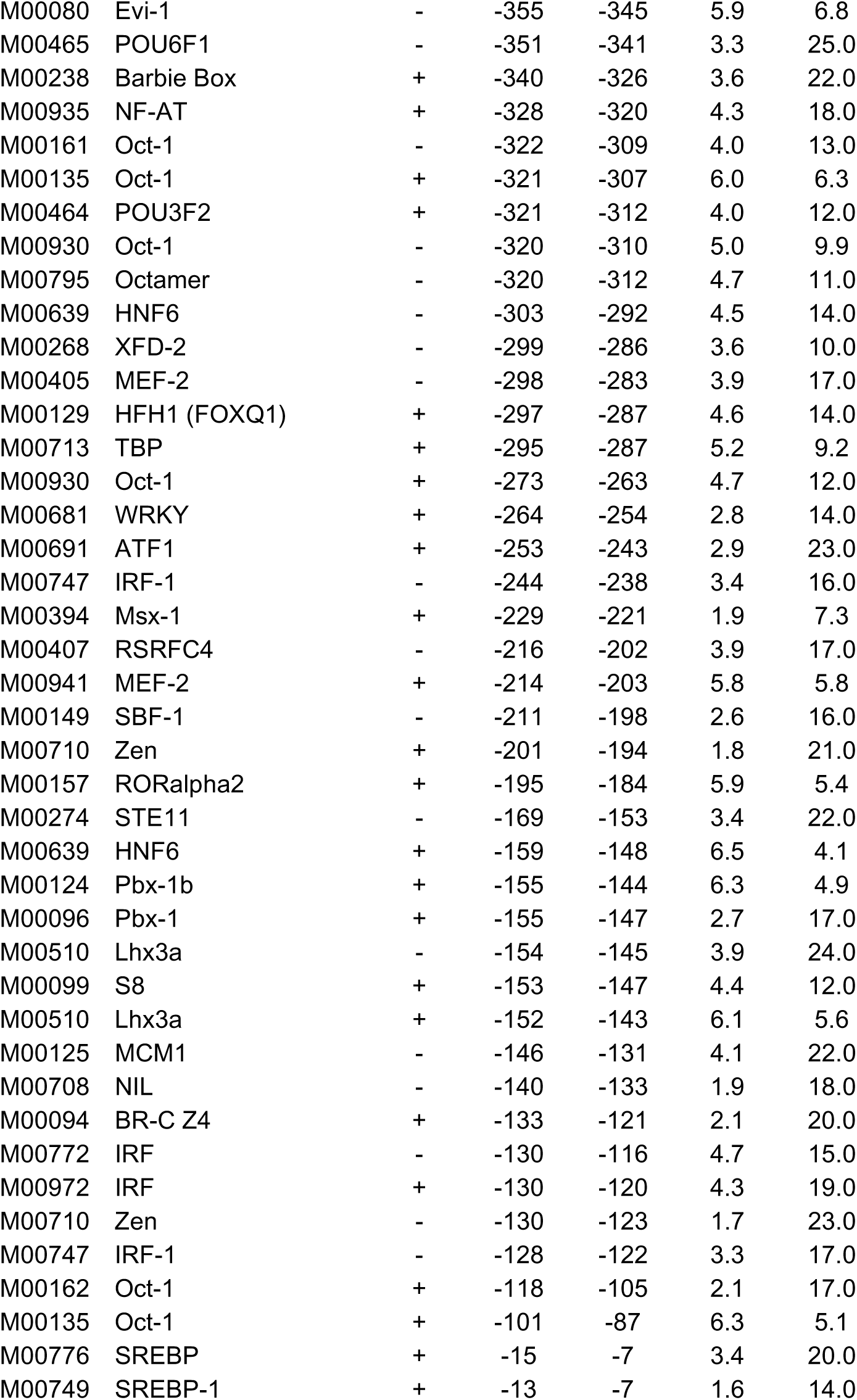
Putative transcription factor-binding sites within the *drosophila dpum* promoter region between -2000 and ±1

**Table EV2.** Putative transcription factor-binding sites within the mouse *pum2* promoter region between -2000 and ±1

| Model | Factor | Strand | Start | End | Score | Eval |
| --- | --- | --- | --- | --- | --- | --- |
| M00057 | COMP1 | + | -1982 | -1960 | 2.8 | 23.0 |
| M00438 | ARF | - | -1972 | -1965 | 2.8 | 9.5 |
| M00135 | Oct-1 | + | -1960 | -1946 | 3.4 | 20.0 |
| M00664 | STE12 | + | -1944 | -1938 | 2.0 | 2.7 |
| M00678 | Tel-2 | - | -1932 | -1923 | 2.6 | 12.0 |
| M00946 | TGA1b | + | -1923 | -1914 | 2.0 | 19.0 |
| M00944 | CPRF-3 | - | -1923 | -1916 | 1.9 | 24.0 |
| M00946 | TGA1b | - | -1923 | -1914 | 2.0 | 19.0 |
| M00788 | EmBP-1b | + | -1922 | -1916 | 3.0 | 6.6 |
| M00472 | FOXO4 | - | -1883 | -1873 | 4.2 | 21.0 |
| M00021 | Kr | - | -1859 | -1850 | 1.8 | 21.0 |
| M00042 | SOX5 | + | -1847 | -1840 | 3.7 | 19.0 |
| M00109 | C/EBPbeta | + | -1845 | -1832 | 3.4 | 25.0 |
| M00621 | C/EBPdelta | - | -1844 | -1833 | 3.4 | 24.0 |
| M00115 | Tax/CREB | - | -1790 | -1776 | 4.4 | 9.9 |
| M00793 | YY1 | - | -1788 | -1780 | 2.6 | 22.0 |
| M00137 | Oct-1 | + | -1772 | -1763 | 3.2 | 19.0 |
| M00423 | FOXJ2 | + | -1772 | -1759 | 3.3 | 9.7 |
| M00267 | XFD-1 | - | -1769 | -1756 | 2.8 | 19.0 |
| M00269 | XFD-3 | - | -1769 | -1756 | 6.0 | 1.8 |
| M00094 | BR-C Z4 | - | -1768 | -1756 | 2.6 | 12.0 |
| M00809 | FOX factors | + | -1768 | -1756 | 5.3 | 8.7 |
| M00962 | AR | + | -1758 | -1750 | 2.8 | 18.0 |
| M00808 | Pax | - | -1738 | -1728 | 3.9 | 5.8 |
| M00060 | Sn | - | -1730 | -1720 | 3.9 | 25.0 |
| M00044 | Sn | + | -1729 | -1716 | 4.7 | 3.5 |
| M00268 | XFD-2 | - | -1727 | -1714 | 3.8 | 8.8 |
| M00632 | GATA-4 | + | -1718 | -1707 | 3.8 | 6.6 |
| M00247 | PacC | - | -1685 | -1669 | 2.6 | 16.0 |
| M00974 | SMAD | + | -1677 | -1667 | 4.4 | 12.0 |
| M00438 | ARF | - | -1667 | -1660 | 2.8 | 9.0 |
| M00457 | STAT5A (homodimer) | - | -1658 | -1645 | 4.5 | 17.0 |
| M00808 | Pax | + | -1652 | -1642 | 4.8 | 2.7 |
| M00723 | GAGA factor | + | -1616 | -1606 | 3.2 | 23.0 |
| M00666 | Sry-beta | + | -1601 | -1593 | 1.8 | 22.0 |
| M00064 | PHO4 | - | -1594 | -1585 | 3.3 | 13.0 |
| M00942 | CPRF-1 | + | -1593 | -1584 | 2.5 | 7.5 |
| M00440 | CG1 | + | -1593 | -1583 | 4.4 | 2.2 |
| M00942 | CPRF-1 | - | -1593 | -1584 | 2.5 | 7.5 |
| M00366 | EmBP-1 | - | -1593 | -1584 | 4.6 | 13.0 |
| M00034 | p53 | - | -1587 | -1568 | 1.4 | 18.0 |
| M00414 | AREB6 | + | -1571 | -1560 | 6.5 | 3.1 |
| M00277 | Lmo2 complex | - | -1571 | -1563 | 4.9 | 13.0 |
| M00060 | Sn | - | -1571 | -1561 | 5.1 | 12.0 |
| M00412 | AREB6 | + | -1570 | -1560 | 3.8 | 18.0 |
| M00413 | AREB6 | + | -1569 | -1560 | 4.2 | 13.0 |
| M00404 | MADS-B | + | -1557 | -1543 | 3.2 | 19.0 |
| M00105 | CDP CR3 | + | -1554 | -1540 | 5.2 | 10.0 |
| M00463 | POU3F2 | + | -1549 | -1538 | 5.0 | 11.0 |
| M00802 | Pit-1 | + | -1549 | -1533 | 5.0 | 11.0 |
| M00162 | Oct-1 | + | -1546 | -1533 | 1.7 | 25.0 |
| M00639 | HNF6 | - | -1545 | -1534 | 3.8 | 21.0 |
| M00405 | MEF-2 | - | -1543 | -1528 | 4.7 | 10.0 |
| M00941 | MEF-2 | - | -1535 | -1524 | 4.2 | 16.0 |
| M00482 | Pitx2 | - | -1525 | -1515 | 4.3 | 5.8 |
| M00763 | PPAR direct repeat 1 | - | -1517 | -1505 | 3.4 | 21.0 |
| M00463 | POU3F2 | + | -1501 | -1488 | 3.9 | 19.0 |
| M00662 | SGF-3 | - | -1500 | -1494 | 2.1 | 13.0 |
| M00033 | p300 | + | -1476 | -1466 | 2.4 | 20.0 |
| M00162 | Oct-1 | - | -1471 | -1458 | 2.8 | 8.5 |
| M00464 | POU3F2 | + | -1470 | -1461 | 3.9 | 12.0 |
| M00407 | RSRFC4 | - | -1469 | -1455 | 3.0 | 25.0 |
| M00406 | MEF-2 | - | -1469 | -1455 | 4.9 | 12.0 |
| M00672 | TEF | - | -1448 | -1437 | 2.3 | 19.0 |
| M00707 | TFIIA | + | -1447 | -1436 | 1.9 | 17.0 |
| M00932 | Sp1 | - | -1432 | -1422 | 4.0 | 24.0 |
| M00931 | Sp1 | - | -1431 | -1422 | 4.1 | 22.0 |
| M00649 | MAZ | - | -1429 | -1422 | 3.4 | 16.0 |
| M00154 | STRE | - | -1426 | -1421 | 2.5 | 23.0 |
| M00404 | MADS-B | - | -1411 | -1397 | 2.9 | 25.0 |
| M00639 | HNF6 | - | -1388 | -1377 | 3.8 | 21.0 |
| M00616 | AFP1 | + | -1386 | -1376 | 3.4 | 7.6 |
| M00026 | RSRFC4 | + | -1357 | -1344 | 6.2 | 5.4 |
| M00967 | HNF4, COUP | - | -1344 | -1337 | 3.4 | 15.0 |
| M00046 | GCR1 | + | -1341 | -1333 | 2.0 | 22.0 |
| M00746 | Elf-1 | - | -1340 | -1329 | 4.0 | 8.7 |
| M00655 | PEA3 | + | -1340 | -1334 | 2.7 | 15.0 |
| M00274 | STE11 | + | -1305 | -1289 | 6.8 | 3.6 |
| M00094 | BR-C Z4 | - | -1304 | -1292 | 3.9 | 3.5 |
| M00802 | Pit-1 | + | -1303 | -1287 | 3.9 | 20.0 |
| M00436 | IPF1 | + | -1298 | -1287 | 3.0 | 13.0 |
| M00937 | TGA1a | + | -1295 | -1274 | 4.6 | 6.6 |
| M00157 | RORalpha2 | - | -1292 | -1281 | 4.1 | 16.0 |
| M00156 | RORalpha1 | - | -1292 | -1282 | 4.5 | 17.0 |
| M00454 | MRF-2 | - | -1282 | -1273 | 2.8 | 24.0 |
| M00354 | Dof3 | + | -1272 | -1252 | 4.1 | 20.0 |
| M00267 | XFD-1 | - | -1249 | -1236 | 2.9 | 18.0 |
| M00809 | FOX factors | + | -1248 | -1236 | 5.3 | 8.9 |
| M00094 | BR-C Z4 | - | -1248 | -1236 | 3.0 | 8.9 |
| M00791 | HNF3 | - | -1247 | -1236 | 4.5 | 15.0 |
| M00724 | HNF3alpha | + | -1245 | -1235 | 5.5 | 8.5 |
| M00639 | HNF6 | - | -1242 | -1231 | 3.7 | 22.0 |
| M00791 | HNF3 | - | -1235 | -1224 | 4.5 | 15.0 |
| M00268 | XFD-2 | - | -1233 | -1220 | 2.7 | 21.0 |
| M00094 | BR-C Z4 | - | -1232 | -1220 | 2.3 | 17.0 |
| M00639 | HNF6 | - | -1230 | -1219 | 4.1 | 18.0 |
| M00724 | HNF3alpha | + | -1229 | -1219 | 5.7 | 7.4 |
| M00062 | IRF-1 | - | -1219 | -1208 | 6.8 | 3.5 |
| M00110 | Elf-1 | + | -1217 | -1204 | 1.3 | 24.0 |
| M00405 | MEF-2 | - | -1214 | -1199 | 4.4 | 12.0 |
| M00734 | CIZ | - | -1213 | -1205 | 4.8 | 7.7 |
| M00979 | Pax-6 | + | -1177 | -1164 | 8.2 | 1.3 |
| M00808 | Pax | + | -1172 | -1162 | 4.9 | 2.3 |
| M00978 | LEF1, TCF1 | + | -1163 | -1153 | 4.0 | 20.0 |
| M00979 | Pax-6 | + | -1148 | -1135 | 6.7 | 3.7 |
| M00727 | SF1 | + | -1147 | -1140 | 3.7 | 13.0 |
| M00005 | AP-4 | - | -1128 | -1111 | 2.8 | 20.0 |
| M00983 | MAF | - | -1118 | -1111 | 3.1 | 13.0 |
| M00480 | LUN-1 | + | -1113 | -1099 | 2.0 | 24.0 |
| M00480 | LUN-1 | - | -1113 | -1099 | 16.6 | 0.0 |
| M00666 | Sry-beta | - | -1108 | -1100 | 1.8 | 21.0 |
| M00623 | Crx | - | -1104 | -1092 | 4.4 | 13.0 |
| M00482 | Pitx2 | - | -1104 | -1094 | 5.3 | 2.6 |
| M00008 | Sp1 | - | -1083 | -1074 | 2.2 | 22.0 |
| M00079 | Evi-1 | + | -1074 | -1065 | 5.2 | 11.0 |
| M00011 | Evi-1 | + | -1073 | -1065 | 4.0 | 13.0 |
| M00133 | Tst-1 | - | -1066 | -1054 | 2.1 | 11.0 |
| M00622 | C/EBPgamma | - | -1065 | -1053 | 4.0 | 7.4 |
| M00701 | SMAD3 | + | -1054 | -1046 | 3.6 | 12.0 |
| M00802 | Pit-1 | - | -1048 | -1032 | 6.8 | 3.5 |
| M00137 | Oct-1 | + | -1043 | -1034 | 3.2 | 19.0 |
| M00820 | HAHB-4 | + | -1042 | -1034 | 5.6 | 8.2 |
| M00480 | LUN-1 | - | -1031 | -1016 | 4.2 | 8.6 |
| M00985 | Stra13 | - | -998 | -988 | 4.3 | 17.0 |
| M00443 | Opaque-2 | - | -996 | -985 | 2.9 | 25.0 |
| M00274 | STE11 | - | -993 | -977 | 4.1 | 16.0 |
| M00978 | LEF1, TCF1 | - | -991 | -981 | 3.9 | 21.0 |
| M00745 | NIL | + | -989 | -980 | 2.9 | 24.0 |
| M00480 | LUN-1 | + | -954 | -938 | 5.7 | 4.3 |
| M00105 | CDP CR3 | + | -953 | -939 | 4.0 | 22.0 |
| M00501 | ANT | + | -953 | -940 | 6.9 | 3.4 |
| M00641 | HSF | - | -949 | -937 | 2.8 | 20.0 |
| M00977 | EBF | - | -947 | -937 | 2.3 | 18.0 |
| M00437 | CHX10 | - | -931 | -921 | 5.0 | 12.0 |
| M00478 | Cdc5 | + | -930 | -919 | 2.2 | 22.0 |
| M00510 | Lhx3a | + | -929 | -920 | 7.4 | 2.3 |
| M00099 | S8 | - | -927 | -921 | 4.2 | 14.0 |
| M00099 | S8 | + | -926 | -920 | 4.2 | 14.0 |
| M00708 | NIL | - | -926 | -919 | 1.7 | 22.0 |
| M00671 | TCF-4 | - | -923 | -916 | 1.8 | 15.0 |
| M00016 | E74A | + | -901 | -887 | 4.1 | 18.0 |
| M00678 | Tel-2 | - | -898 | -889 | 3.5 | 4.9 |
| M00023 | HOXA5 (Hox-1.3) | - | -896 | -867 | 6.3 | 4.6 |
| M00018 | Ubx | - | -890 | -882 | 5.3 | 10.0 |
| M00489 | Nkx6-2 | + | -886 | -876 | 1.5 | 22.0 |
| M00510 | Lhx3a | + | -886 | -877 | 5.9 | 6.8 |
| M00099 | S8 | - | -884 | -878 | 4.2 | 14.0 |
| M00437 | CHX10 | + | -883 | -873 | 3.8 | 23.0 |
| M00099 | S8 | + | -883 | -877 | 4.3 | 14.0 |
| M00016 | E74A | + | -882 | -868 | 5.5 | 8.3 |
| M00655 | PEA3 | - | -878 | -872 | 2.7 | 15.0 |
| M00634 | GCM | - | -866 | -855 | 2.6 | 19.0 |
| M00410 | SOX9 | - | -796 | -787 | 5.5 | 8.9 |
| M00728 | ROX1 | + | -795 | -787 | 4.4 | 16.0 |
| M00978 | LEF1, TCF1 | + | -794 | -784 | 3.9 | 21.0 |
| M00042 | SOX5 | - | -794 | -787 | 3.7 | 18.0 |
| M00079 | Evi-1 | + | -763 | -754 | 4.6 | 15.0 |
| M00790 | HNF1 | - | -746 | -729 | 3.7 | 25.0 |
| M00269 | XFD-3 | + | -738 | -725 | 3.1 | 18.0 |
| M00809 | FOX factors | - | -738 | -726 | 5.1 | 9.9 |
| M00130 | FOXD3 | - | -737 | -727 | 5.3 | 9.8 |
| M00772 | IRF | + | -707 | -693 | 4.3 | 18.0 |
| M00699 | ICSBP | - | -705 | -694 | 4.2 | 10.0 |
| M00984 | PEBP | - | -700 | -686 | 3.5 | 23.0 |
| M00711 | Zta | + | -682 | -670 | 4.2 | 10.0 |
| M00459 | STAT5B (homodimer) | + | -667 | -654 | 3.5 | 23.0 |
| M00457 | STAT5A (homodimer) | + | -667 | -654 | 5.9 | 6.4 |
| M00746 | Elf-1 | + | -666 | -655 | 3.7 | 12.0 |
| M00007 | Elk-1 | + | -665 | -651 | 2.3 | 14.0 |
| M00531 | NERF1a | + | -665 | -648 | 4.9 | 6.7 |
| M00108 | NRF-2/GABP | + | -663 | -654 | 2.2 | 25.0 |
| M00046 | GCR1 | - | -662 | -654 | 2.0 | 22.0 |
| M00743 | c-Ets-1 68 | + | -662 | -655 | 2.9 | 15.0 |
| M00678 | Tel-2 | - | -662 | -653 | 3.9 | 3.3 |
| M00032 | c-Ets-1(p54) | + | -662 | -654 | 3.7 | 15.0 |
| M00771 | Ets | - | -662 | -655 | 5.7 | 7.5 |
| M00655 | PEA3 | - | -661 | -655 | 2.7 | 15.0 |
| M00515 | PPARgamma:RXRalpha, PPARgamma | - | -658 | -636 | 2.8 | 12.0 |
| M00447 | AR | - | -629 | -615 | 2.8 | 25.0 |
| M00753 | RCS1 | - | -627 | -615 | 3.9 | 12.0 |
| M00408 | MADS-A | - | -622 | -607 | 2.7 | 16.0 |
| M00081 | Evi-1 | - | -617 | -603 | 3.3 | 12.0 |
| M00022 | Hb | - | -612 | -603 | 4.0 | 13.0 |
| M00934 | Zeste | + | -589 | -580 | 6.8 | 3.6 |
| M00444 | VDR | + | -583 | -569 | 2.9 | 23.0 |
| M00154 | STRE | + | -576 | -571 | 2.5 | 23.0 |
| M00941 | MEF-2 | + | -551 | -540 | 3.9 | 18.0 |
| M00016 | E74A | + | -514 | -500 | 6.8 | 3.6 |
| M00108 | NRF-2/GABP | + | -512 | -503 | 3.0 | 13.0 |
| M00743 | c-Ets-1 68 | + | -511 | -504 | 3.3 | 11.0 |
| M00032 | c-Ets-1(p54) | + | -511 | -503 | 4.0 | 12.0 |
| M00678 | Tel-2 | - | -511 | -502 | 4.2 | 2.3 |
| M00025 | Elk-1 | + | -511 | -502 | 5.2 | 11.0 |
| M00655 | PEA3 | - | -510 | -504 | 2.4 | 21.0 |
| M00084 | MZF1 | + | -508 | -496 | 4.6 | 7.5 |
| M00404 | MADS-B | - | -490 | -476 | 3.3 | 18.0 |
| M00633 | GBF | + | -483 | -475 | 2.5 | 7.7 |
| M00749 | SREBP-1 | - | -481 | -475 | 1.7 | 13.0 |
| M00670 | NIL | + | -450 | -443 | 1.8 | 9.2 |
| M00141 | Lyf-1 | + | -444 | -436 | 2.6 | 21.0 |
| M00275 | Mat1-Mc | - | -441 | -424 | 3.6 | 18.0 |
| M00162 | Oct-1 | + | -436 | -423 | 1.8 | 24.0 |
| M00728 | ROX1 | - | -436 | -428 | 4.7 | 13.0 |
| M00042 | SOX5 | + | -436 | -429 | 3.7 | 18.0 |
| M00410 | SOX9 | + | -436 | -427 | 5.5 | 8.8 |
| M00708 | NIL | - | -434 | -427 | 1.9 | 18.0 |
| M00632 | GATA-4 | + | -429 | -418 | 3.7 | 7.3 |
| M00531 | NERF1a | + | -422 | -405 | 3.4 | 19 |
| M00771 | Ets | - | -419 | -412 | 4 | 24 |
| M00681 | WRKY | - | -409 | -399 | 2.3 | 20 |
| M00744 | POU1F1 | + | -406 | -397 | 6.2 | 4.5 |
| M00463 | POU3F2 | + | -406 | -393 | 6.5 | 4.4 |
| M00463 | POU3F2 | - | -406 | -393 | 8.8 | 0.88 |
| M00662 | SGF-3 | - | -405 | -399 | 1.6 | 21 |
| M00274 | STE11 | - | -394 | -378 | 3.5 | 21 |
| M00670 | NIL | + | -376 | -369 | 0.9 | 23 |
| M00115 | Tax/CREB | - | -376 | -362 | 4.8 | 7.5 |
| M00479 | Alfin1 | - | -367 | -357 | 2.5 | 23 |
| M00751 | AML1 | - | -363 | -358 | 2.1 | 9.8 |
| M00237 | AhR:Arnt | + | -358 | -340 | 5.9 | 6.7 |
| M00448 | Zic1 | - | -347 | -339 | 2.2 | 22 |
| M00650 | MTF-1 | - | -342 | -329 | 4.2 | 17 |
| M00723 | GAGA factor | + | -333 | -323 | 3.4 | 19 |
| M00158 | COUP-TF, HNF-4 | - | -329 | -316 | 3.8 | 22 |
| M00765 | COUP direct repeat 1 | - | -328 | -316 | 3.3 | 25 |
| M00763 | PPAR direct repeat 1 | - | -328 | -316 | 3.4 | 21 |
| M00629 | Eve | - | -310 | -301 | 1.6 | 25 |
| M00720 | CAC-binding protein | - | -306 | -298 | 2.7 | 17 |
| M00777 | STAT | - | -295 | -288 | 5.2 | 10 |
| M00249 | CHOP:C/EBPalpha | - | -277 | -267 | 4 | 22 |
| M00069 | YY1 | - | -271 | -255 | 5.3 | 6.6 |
| M00733 | SMAD4 | - | -263 | -249 | 3.2 | 21 |
| M00651 | NF-muE1 | - | -263 | -255 | 4.2 | 5.5 |
| M00652 | NRF-1 | - | -225 | -216 | 6.8 | 2.4 |
| M00778 | AhR | - | -224 | -215 | 3.7 | 23 |
| M00652 | NRF-1 | + | -223 | -214 | 3.5 | 19 |
| M00235 | AhR:Arnt | - | -222 | -215 | 3.9 | 18 |
| M00650 | MTF-1 | + | -197 | -184 | 5.7 | 7.5 |
| M00716 | ZF5 | + | -187 | -180 | 3.1 | 11 |
| M00433 | Hmx3 (Nkx5-1) | - | -173 | -164 | 3.1 | 21 |
| M00247 | PacC | + | -164 | -148 | 2.5 | 19 |
| M00644 | LBP-1 | + | -149 | -143 | 1.1 | 13 |
| M00632 | GATA-4 | - | -134 | -123 | 3.2 | 11 |
| M00800 | AP-2 | - | -115 | -100 | 3.4 | 22 |
| M00650 | MTF-1 | + | -112 | -99 | 6.8 | 3.6 |
| M00333 | ZF5 | - | -111 | -100 | 2.6 | 15 |
| M00716 | ZF5 | + | -109 | -102 | 3.1 | 11 |
| M00721 | CACCC-binding factor | + | -104 | -89 | 4.7 | 5.9 |
| M00915 | AP-2 | - | -103 | -91 | 4 | 18 |
| M00800 | AP-2 | + | -102 | -87 | 3.7 | 19 |
| M00933 | Sp1 | - | -98 | -89 | 5.1 | 11 |
| M00807 | Egr | + | -98 | -88 | 3 | 15 |
| M00938 | E2F-1 | + | -97 | -82 | 3.3 | 19 |
| M00982 | KROX | - | -97 | -84 | 3.7 | 24 |
| M00932 | Sp1 | + | -97 | -87 | 4.7 | 15 |
| M00444 | VDR | + | -92 | -78 | 3.2 | 18 |
| M00649 | MAZ | + | -91 | -84 | 3.4 | 16 |
| M00706 | TFII-I | + | -89 | -81 | 1.9 | 12 |
| M00154 | STRE | + | -83 | -78 | 2.5 | 23 |
| M00649 | MAZ | + | -82 | -75 | 3.4 | 16 |
| M00982 | KROX | - | -80 | -67 | 3.9 | 21 |
| M00923 | Adf-1 | - | -76 | -56 | 8.3 | 1.2 |
| M00695 | ETF | + | -71 | -65 | 2.8 | 21 |
| M00069 | YY1 | + | -61 | -45 | 3.3 | 24 |
| M00793 | YY1 | + | -59 | -51 | 3.9 | 8.3 |
| M00808 | Pax | - | -48 | -38 | 2.2 | 25 |
| M00721 | CACCC-binding factor | - | -35 | -20 | 2.7 | 25 |

**Table EV3.** Putative transcription factor-binding sites within the human *pum2* promoter region between -2000 and ±1

| Model | Factor | Strand | Start | End | Score | Eval |
| --- | --- | --- | --- | --- | --- | --- |
| M00506 | LIM1 | + | -1996 | -1985 | 2.5 | 14.0 |
| M00753 | RCS1 | + | -1994 | -1982 | 3.4 | 17.0 |
| M00407 | RSRFC4 | - | -1978 | -1964 | 4.3 | 15.0 |
| M00408 | MADS-A | - | -1977 | -1962 | 2.9 | 14.0 |
| M00404 | MADS-B | - | -1977 | -1963 | 5.9 | 3.0 |
| M00026 | RSRFC4 | + | -1975 | -1962 | 4.7 | 12.0 |
| M00666 | Sry-beta | - | -1963 | -1955 | 1.8 | 22.0 |
| M00054 | NF-kappaB | + | -1957 | -1948 | 3.9 | 22.0 |
| M00143 | Pax-5 | - | -1937 | -1910 | 3.8 | 17.0 |
| M00683 | XBP1 | - | -1929 | -1924 | 2.8 | 14.0 |
| M00518 | PPARalpha:RXRalpha | - | -1929 | -1915 | 5.6 | 8.3 |
| M00155 | ARP-1 (COUP-TF2) | + | -1927 | -1912 | 3.9 | 6.7 |
| M00963 | T3R | + | -1921 | -1913 | 3.5 | 17.0 |
| M00647 | LXRalpha:RXRalpha | - | -1919 | -1902 | 5.8 | 5.6 |
| M00766 | LXR direct repeat 4 | + | -1919 | -1904 | 9.2 | 0.6 |
| M00733 | SMAD4 | - | -1912 | -1898 | 3.4 | 18.0 |
| M00932 | Sp1 | - | -1909 | -1899 | 4.1 | 22.0 |
| M00807 | Egr | - | -1908 | -1898 | 2.5 | 23.0 |
| M00931 | Sp1 | - | -1908 | -1899 | 4.4 | 19.0 |
| M00933 | Sp1 | + | -1907 | -1898 | 4.5 | 17.0 |
| M00982 | KROX | + | -1906 | -1893 | 4.1 | 19.0 |
| M00700 | ROM | + | -1902 | -1894 | 3.2 | 6.4 |
| M00506 | LIM1 | + | -1901 | -1890 | 2.3 | 17.0 |
| M00141 | Lyf-1 | - | -1892 | -1884 | 4.9 | 3.3 |
| M00480 | LUN-1 | - | -1890 | -1875 | 19.7 | 0.0 |
| M00666 | Sry-beta | - | -1884 | -1876 | 1.8 | 21.0 |
| M00482 | Pitx2 | - | -1880 | -1870 | 4.8 | 4.1 |
| M00717 | Pax-8 | + | -1868 | -1861 | 1.5 | 20.0 |
| M00725 | HP1 site factor | - | -1838 | -1827 | 3.1 | 13.0 |
| M00006 | MEF-2 | + | -1837 | -1822 | 3.9 | 15.0 |
| M00406 | MEF-2 | + | -1836 | -1822 | 6.3 | 5.1 |
| M00753 | RCS1 | + | -1831 | -1819 | 3.2 | 20.0 |
| M00018 | Ubx | - | -1803 | -1795 | 4.2 | 20.0 |
| M00478 | Cdc5 | + | -1766 | -1755 | 3.3 | 7.7 |
| M00465 | POU6F1 | - | -1764 | -1754 | 3.8 | 18.0 |
| M00464 | POU3F2 | - | -1764 | -1755 | 3.9 | 13.0 |
| M00689 | unc-86 | - | -1756 | -1747 | 2.2 | 16.0 |
| M00725 | HP1 site factor | + | -1750 | -1739 | 3.7 | 8.3 |
| M00005 | AP-4 | - | -1748 | -1731 | 2.6 | 24.0 |
| M00626 | RFX1 (EF-C) | - | -1746 | -1733 | 2.4 | 18.0 |
| M00485 | Nkx2-2 | - | -1710 | -1701 | 4.2 | 18.0 |
| M00105 | CDP CR3 | - | -1668 | -1654 | 4.1 | 20.0 |
| M00935 | NF-AT | + | -1655 | -1647 | 4.5 | 16.0 |
| M00749 | SREBP-1 | + | -1604 | -1598 | 2.0 | 9.5 |
| M00154 | STRE | + | -1577 | -1572 | 2.5 | 23.0 |
| M00649 | MAZ | + | -1576 | -1569 | 3.4 | 16.0 |
| M00960 | PR, GR | + | -1571 | -1562 | 4.6 | 11.0 |
| M00930 | Oct-1 | + | -1541 | -1531 | 3.5 | 24.0 |
| M00977 | EBF | + | -1537 | -1527 | 2.0 | 23.0 |
| M00769 | AML | + | -1533 | -1522 | 3.9 | 21.0 |
| M00751 | AML1 | + | -1531 | -1526 | 2.1 | 9.8 |
| M00722 | core-binding factor | + | -1531 | -1524 | 3.2 | 12.0 |
| M00501 | ANT | - | -1511 | -1498 | 7.7 | 1.9 |
| M00975 | RFX | - | -1499 | -1491 | 2.6 | 14.0 |
| M00093 | BR-C Z3 | - | -1489 | -1478 | 2.9 | 15.0 |
| M00011 | Evi-1 | - | -1489 | -1481 | 3.7 | 16.0 |
| M00268 | XFD-2 | - | -1487 | -1474 | 3.3 | 12.0 |
| M00472 | FOXO4 | - | -1487 | -1477 | 4.0 | 24.0 |
| M00269 | XFD-3 | - | -1487 | -1474 | 5.3 | 3.2 |
| M00474 | FOXO1 | + | -1486 | -1474 | 9.0 | 0.8 |
| M00619 | Alx-4 | - | -1482 | -1470 | 4.0 | 7.3 |
| M00977 | EBF | - | -1476 | -1466 | 2.9 | 10.0 |
| M00154 | STRE | + | -1472 | -1467 | 2.5 | 23.0 |
| M00649 | MAZ | + | -1471 | -1464 | 3.4 | 16.0 |
| M00355 | PBF | - | -1454 | -1445 | 3.4 | 22.0 |
| M00275 | Mat1-Mc | + | -1423 | -1406 | 3.3 | 22.0 |
| M00059 | YY1 | - | -1403 | -1392 | 2.5 | 21.0 |
| M00487 | mtTFA | - | -1395 | -1387 | 2.9 | 16.0 |
| M00622 | C/EBPgamma | + | -1381 | -1369 | 4.0 | 7.3 |
| M00045 | E4BP4 | - | -1379 | -1369 | 4.3 | 17.0 |
| M00616 | AFP1 | - | -1350 | -1340 | 2.6 | 16.0 |
| M00089 | Athb-1 | + | -1346 | -1337 | 6.4 | 4.8 |
| M00775 | NF-Y | - | -1338 | -1328 | 6.1 | 5.9 |
| M00094 | BR-C Z4 | - | -1326 | -1314 | 2.3 | 16.0 |
| M00423 | FOXJ2 | - | -1317 | -1304 | 2.2 | 25.0 |
| M00626 | RFX1 (EF-C) | - | -1306 | -1293 | 2.2 | 22.0 |
| M00053 | c-Rel | - | -1298 | -1289 | 3.5 | 19.0 |
| M00034 | p53 | + | -1298 | -1279 | 5.6 | 4.0 |
| M00034 | p53 | - | -1298 | -1279 | 5.0 | 5.0 |
| M00011 | Evi-1 | - | -1286 | -1278 | 5.1 | 7.2 |
| M00161 | Oct-1 | - | -1276 | -1263 | 4.6 | 8.7 |
| M00795 | Octamer | - | -1274 | -1266 | 3.4 | 25.0 |
| M00020 | Ftz | + | -1267 | -1256 | 3.2 | 25.0 |
| M00708 | NIL | + | -1265 | -1258 | 2.2 | 13.0 |
| M00806 | NF-1 | + | -1240 | -1234 | 3.4 | 23.0 |
| M00261 | Olf-1 | - | -1220 | -1201 | 4.6 | 11.0 |
| M00977 | EBF | - | -1214 | -1204 | 2.6 | 14.0 |
| M00749 | SREBP-1 | - | -1191 | -1185 | 1.7 | 13.0 |
| M00133 | Tst-1 | + | -1184 | -1172 | 1.6 | 19.0 |
| M00267 | XFD-1 | - | -1149 | -1136 | 3.6 | 10.0 |
| M00135 | Oct-1 | - | -1149 | -1135 | 3.9 | 17.0 |
| M00672 | TEF | + | -1146 | -1135 | 2.1 | 22.0 |
| M00930 | Oct-1 | + | -1146 | -1136 | 4.0 | 18.0 |
| M00724 | HNF3alpha | + | -1145 | -1135 | 4.3 | 19.0 |
| M00795 | Octamer | + | -1144 | -1136 | 4.0 | 17.0 |
| M00010 | Opaque-2 | + | -1119 | -1104 | 5.0 | 5.2 |
| M00405 | MEF-2 | - | -1108 | -1093 | 4.3 | 14.0 |
| M00405 | MEF-2 | - | -1091 | -1076 | 4.2 | 15.0 |
| M00696 | En | + | -1089 | -1083 | 3.0 | 12.0 |
| M00640 | HOXA4 | - | -1089 | -1082 | 2.1 | 8.8 |
| M00099 | S8 | + | -1088 | -1082 | 3.8 | 18.0 |
| M00268 | XFD-2 | + | -1086 | -1073 | 2.5 | 24.0 |
| M00708 | NIL | - | -1082 | -1075 | 2.2 | 13.0 |
| M00501 | ANT | - | -1005 | -992 | 5.4 | 9.6 |
| M00444 | VDR | + | -984 | -970 | 3.1 | 19.0 |
| M00721 | CACCC-binding factor | + | -960 | -946 | 2.8 | 24.0 |
| M00720 | CAC-binding protein | + | -955 | -947 | 2.5 | 20.0 |
| M00931 | Sp1 | + | -954 | -945 | 4.5 | 17.0 |
| M00932 | Sp1 | + | -954 | -944 | 4.9 | 13.0 |
| M00479 | Alfin1 | + | -953 | -943 | 4.0 | 6.5 |
| M00684 | XPF-1 | + | -929 | -920 | 1.3 | 17.0 |
| M00158 | COUP-TF, HNF-4 | - | -889 | -876 | 3.7 | 22.0 |
| M00161 | Oct-1 | + | -843 | -830 | 3.2 | 22.0 |
| M00269 | XFD-3 | - | -837 | -824 | 2.7 | 24.0 |
| M00268 | XFD-2 | - | -837 | -824 | 3.2 | 14.0 |
| M00463 | POU3F2 | - | -832 | -820 | 3.7 | 22.0 |
| M00408 | MADS-A | - | -826 | -811 | 2.2 | 25.0 |
| M00744 | POU1F1 | + | -825 | -816 | 3.5 | 24.0 |
| M00463 | POU3F2 | + | -825 | -812 | 6.1 | 5.8 |
| M00162 | Oct-1 | + | -822 | -809 | 3.0 | 7.4 |
| M00639 | HNF6 | - | -819 | -808 | 3.9 | 20.0 |
| M00105 | CDP CR3 | + | -803 | -789 | 3.8 | 23.0 |
| M00006 | MEF-2 | + | -790 | -775 | 3.3 | 22.0 |
| M00129 | HFH1 (FOXQ1) | - | -784 | -774 | 4.8 | 13.0 |
| M00438 | ARF | + | -774 | -767 | 2.2 | 17.0 |
| M00484 | Ncx | + | -746 | -739 | 1.9 | 17.0 |
| M00622 | C/EBPgamma | + | -733 | -721 | 4.3 | 6.0 |
| M00772 | IRF | - | -726 | -712 | 6.1 | 5.6 |
| M00699 | ICSBP | + | -725 | -714 | 5.4 | 4.5 |
| M00162 | Oct-1 | - | -725 | -712 | 2.5 | 12.0 |
| M00161 | Oct-1 | - | -725 | -712 | 6.2 | 2.9 |
| M00747 | IRF-1 | - | -724 | -718 | 3.3 | 17.0 |
| M00795 | Octamer | - | -723 | -715 | 3.6 | 22.0 |
| M00930 | Oct-1 | - | -723 | -713 | 5.8 | 6.1 |
| M00154 | STRE | + | -713 | -708 | 2.3 | 25.0 |
| M00734 | CIZ | + | -699 | -691 | 3.9 | 14.0 |
| M00059 | YY1 | - | -684 | -673 | 2.3 | 25.0 |
| M00036 | v-Jun | + | -675 | -661 | 3.7 | 18.0 |
| M00916 | CREB | + | -674 | -661 | 4.8 | 14.0 |
| M00691 | ATF1 | - | -672 | -662 | 3.8 | 12.0 |
| M00917 | CREB | - | -672 | -662 | 4.1 | 23.0 |
| M00775 | NF-Y | + | -658 | -648 | 5.1 | 11.0 |
| M00022 | Hb | - | -641 | -632 | 3.3 | 20.0 |
| M00915 | AP-2 | + | -632 | -620 | 7.3 | 2.4 |
| M00800 | AP-2 | + | -630 | -615 | 3.9 | 16.0 |
| M00158 | COUP-TF, HNF-4 | - | -613 | -600 | 4.4 | 15.0 |
| M00985 | Stra13 | - | -605 | -595 | 7.0 | 3.0 |
| M00440 | CG1 | - | -604 | -594 | 2.6 | 11.0 |
| M00796 | USF | - | -604 | -593 | 6.4 | 4.3 |
| M00942 | CPRF-1 | + | -603 | -594 | 1.3 | 23.0 |
| M00942 | CPRF-1 | - | -603 | -594 | 1.3 | 23.0 |
| M00946 | TGA1b | + | -603 | -594 | 2.1 | 17.0 |
| M00946 | TGA1b | - | -603 | -594 | 2.1 | 17.0 |
| M00440 | CG1 | + | -603 | -593 | 2.8 | 8.9 |
| M00944 | CPRF-3 | - | -603 | -596 | 2.5 | 14.0 |
| M00945 | CPRF-2 | - | -603 | -596 | 3.1 | 21.0 |
| M00366 | EmBP-1 | - | -603 | -594 | 4.2 | 17.0 |
| M00788 | EmBP-1b | + | -602 | -596 | 3.4 | 3.8 |
| M00985 | Stra13 | + | -602 | -592 | 5.2 | 11.0 |
| M00727 | SF1 | + | -582 | -575 | 3.7 | 13.0 |
| M00710 | Zen | - | -553 | -546 | 1.6 | 25.0 |
| M00463 | POU3F2 | - | -549 | -536 | 3.4 | 25.0 |
| M00091 | BR-C Z1 | + | -507 | -490 | 2.9 | 21.0 |
| M00274 | STE11 | - | -504 | -488 | 3.6 | 20.0 |
| M00275 | Mat1-Mc | - | -504 | -487 | 5.8 | 4.0 |
| M00724 | HNF3alpha | - | -502 | -492 | 4.8 | 14.0 |
| M00105 | CDP CR3 | + | -499 | -485 | 3.6 | 25.0 |
| M00267 | XFD-1 | + | -497 | -484 | 3.1 | 15.0 |
| M00488 | DREF | - | -496 | -483 | 3.4 | 22.0 |
| M00941 | MEF-2 | - | -494 | -483 | 3.5 | 24.0 |
| M00354 | Dof3 | - | -487 | -467 | 4.1 | 19.0 |
| M00441 | GBF | + | -483 | -472 | 4.9 | 8.6 |
| M00440 | CG1 | + | -482 | -472 | 2.3 | 14.0 |
| M00942 | CPRF-1 | + | -482 | -473 | 3.7 | 2.4 |
| M00366 | EmBP-1 | + | -482 | -473 | 4.5 | 14.0 |
| M00942 | CPRF-1 | - | -482 | -473 | 3.7 | 2.4 |
| M00366 | EmBP-1 | - | -482 | -473 | 7.1 | 2.9 |
| M00788 | EmBP-1b | + | -481 | -475 | 3.4 | 4.2 |
| M00943 | TAF-1 | + | -480 | -473 | 2.0 | 21.0 |
| M00944 | CPRF-3 | + | -480 | -473 | 2.1 | 20.0 |
| M00945 | CPRF-2 | + | -480 | -473 | 3.1 | 21.0 |
| M00972 | IRF | + | -452 | -442 | 4.1 | 22.0 |
| M00747 | IRF-1 | + | -442 | -436 | 3.4 | 16.0 |
| M00116 | C/EBPalpha | - | -430 | -420 | 3.3 | 19.0 |
| M00159 | C/EBP | - | -429 | -419 | 2.1 | 25.0 |
| M00135 | Oct-1 | - | -414 | -400 | 4.4 | 14.0 |
| M00223 | STATx | + | -408 | -400 | 3.7 | 21.0 |
| M00027 | AbaA | - | -398 | -383 | 4.7 | 15.0 |
| M00777 | STAT | - | -396 | -389 | 4.0 | 23.0 |
| M00701 | SMAD3 | - | -365 | -357 | 3.5 | 13.0 |
| M00733 | SMAD4 | + | -364 | -350 | 3.3 | 19.0 |
| M00247 | PacC | + | -360 | -344 | 3.4 | 8.1 |
| M00804 | E2A | - | -349 | -337 | 3.5 | 17.0 |
| M00031 | MATalpha2 | + | -342 | -334 | 2.6 | 7.8 |
| M00974 | SMAD | - | -254 | -244 | 4.3 | 13.0 |
| M00480 | LUN-1 | + | -238 | -223 | 3.6 | 12.0 |
| M00967 | HNF4, COUP | + | -234 | -227 | 3.7 | 12.0 |
| M00683 | XBP1 | + | -220 | -215 | 2.8 | 14.0 |
| M00482 | Pitx2 | + | -204 | -194 | 3.0 | 16.0 |
| M00059 | YY1 | + | -191 | -180 | 2.8 | 15.0 |
| M00649 | MAZ | + | -181 | -174 | 3.1 | 20.0 |
| M00931 | Sp1 | + | -181 | -172 | 3.9 | 25.0 |
| M00639 | HNF6 | - | -169 | -158 | 4.5 | 14.0 |
| M00011 | Evi-1 | - | -161 | -153 | 6.4 | 3.3 |
| M00751 | AML1 | - | -148 | -143 | 2.1 | 10.0 |
| M00749 | SREBP-1 | + | -146 | -140 | 1.6 | 14.0 |
| M00405 | MEF-2 | + | -134 | -119 | 4 | 16 |
| M00713 | TBP | - | -130 | -122 | 5.2 | 9.2 |
| M00407 | RSRFC4 | - | -117 | -103 | 2.9 | 25 |
| M00026 | RSRFC4 | + | -114 | -101 | 5.6 | 8.3 |
| M00154 | STRE | + | -95 | -90 | 2.5 | 23 |
| M00749 | SREBP-1 | + | -64 | -58 | 1.7 | 13 |
| M00622 | C/EBPgamma | - | -26 | -14 | 2.6 | 23 |
| M00707 | TFIIA | + | -24 | -13 | 2.6 | 8.3 |

## References

Akimaru H, Chen Y, Dai P, Hou DX, Nonaka M, Smolik SM, Armstrong S, Goodman RH, Ishii S (1997a) Drosophila CBP is a co-activator of cubitus interruptus in hedgehog signalling. Nature 386: 735-8

Akimaru H, Hou DX, Ishii S (1997b) Drosophila CBP is required for dorsal-dependent twist gene expression. Nat Genet 17: 211-4

Arvola RM, Weidmann CA, Tanaka Hall TM, Goldstrohm AC (2017) Combinatorial control of messenger RNAs by Pumilio, Nanos and Brain Tumor Proteins. RNA Biol: 1-12

Bohn JA, Van Etten JL, Schagat TL, Bowman BM, McEachin RC, Freddolino PL, Goldstrohm AC (2018) Identification of diverse target RNAs that are functionally regulated by human Pumilio proteins. Nucleic Acids Res 46: 362-386

Carreira-Rosario A, Bhargava V, Hillebrand J, Kollipara RK, Ramaswami M, Buszczak M (2016) Repression of Pumilio Protein Expression by Rbfox1 Promotes Germ Cell Differentiation. Dev Cell 36: 562-71

Chen G, Li W, Zhang QS, Regulski M, Sinha N, Barditch J, Tully T, Krainer AR, Zhang MQ, Dubnau J (2008) Identification of synaptic targets of Drosophila pumilio. PLoS Comput Biol 4: e1000026

Chen H, Hung MC (1997) Involvement of co-activator p300 in the transcriptional regulation of the HER-2/neu gene. J Biol Chem 272: 6101-4

Cudmore RH, Turrigiano GG (2004) Long-term potentiation of intrinsic excitability in LV visual cortical neurons. J Neurophysiol 92: 341-8

De Luca A, Severino A, De Paolis P, Cottone G, De Luca L, De Falco M, Porcellini A, Volpe M, Condorelli G (2003) p300/cAMP-response-element-binding-protein (’CREB’)-binding protein (CBP) modulates co-operation between myocyte enhancer factor 2A (MEF2A) and thyroid hormone receptor-retinoid X receptor. Biochem J 369: 477-84

Driscoll HE, Muraro NI, He M, Baines RA (2013) Pumilio-2 regulates translation of Nav1.6 to mediate homeostasis of membrane excitability. J Neurosci 33: 9644-54

Fiore R, Khudayberdiev S, Christensen M, Siegel G, Flavell SW, Kim TK, Greenberg ME, Schratt G (2009) Mef2-mediated transcription of the miR379-410 cluster regulates activity-dependent dendritogenesis by fine-tuning Pumilio2 protein levels. EMBO J 28: 697-710

Fiore R, Rajman M, Schwale C, Bicker S, Antoniou A, Bruehl C, Draguhn A, Schratt G (2014) MiR-134-dependent regulation of Pumilio-2 is necessary for homeostatic synaptic depression. EMBO J 33: 2231-46

Flavell SW, Cowan CW, Kim TK, Greer PL, Lin Y, Paradis S, Griffith EC, Hu LS, Chen C, Greenberg ME (2006) Activity-dependent regulation of MEF2 transcription factors suppresses excitatory synapse number. Science 311: 1008-12

Follwaczny P, Schieweck R, Riedemann T, Demleitner A, Straub T, Klemm AH, Bilban M, Sutor B, Popper B, Kiebler MA (2017) Pumilio2-deficient mice show a predisposition for epilepsy. Dis Model Mech 10: 1333-1342

Gerber AP, Luschnig S, Krasnow MA, Brown PO, Herschlag D (2006) Genome-wide identification of mRNAs associated with the translational regulator PUMILIO in Drosophila melanogaster. Proc Natl Acad Sci U S A 103: 4487-92

Giachello CN, Baines RA (2017) Regulation of motoneuron excitability and the setting of homeostatic limits. Curr Opin Neurobiol 43: 1-6

Gossett LA, Kelvin DJ, Sternberg EA, Olson EN (1989) A new myocyte-specific enhancer-binding factor that recognizes a conserved element associated with multiple muscle-specific genes. Mol Cell Biol 9: 5022-33

Gunay C, Prinz AA (2010) Model calcium sensors for network homeostasis: sensor and readout parameter analysis from a database of model neuronal networks. J Neurosci 30: 1686-98

Gunthorpe D, Beatty KE, Taylor MV (1999) Different levels, but not different isoforms, of the Drosophila transcription factor DMEF2 affect distinct aspects of muscle differentiation. Dev Biol 215: 130-45

Han J, Jiang Y, Li Z, Kravchenko VV, Ulevitch RJ (1997) Activation of the transcription factor MEF2C by the MAP kinase p38 in inflammation. Nature 386: 296-9

Hansen KF, Sakamoto K, Pelz C, Impey S, Obrietan K (2014) Profiling status epilepticus-induced changes in hippocampal RNA expression using high-throughput RNA sequencing. Sci Rep 4: 6930

Kao HY, Verdel A, Tsai CC, Simon C, Juguilon H, Khochbin S (2001) Mechanism for nucleocytoplasmic shuttling of histone deacetylase 7. J Biol Chem 276: 47496-507

Lachmann A, Xu H, Krishnan J, Berger SI, Mazloom AR, Ma’ayan A (2010) ChEA: transcription factor regulation inferred from integrating genome-wide ChIP-X experiments. Bioinformatics 26: 2438-44

Lilly B, Galewsky S, Firulli AB, Schulz RA, Olson EN (1994) D-MEF2: a MADS box transcription factor expressed in differentiating mesoderm and muscle cell lineages during Drosophila embryogenesis. Proc Natl Acad Sci U S A 91: 5662-6

Lin WH, Giachello CN, Baines RA (2017) Seizure control through genetic and pharmacological manipulation of Pumilio in Drosophila: a key component of neuronal homeostasis. Dis Model Mech 10: 141-150

Lin WH, He M, Baines RA (2015) Seizure suppression through manipulating splicing of a voltage-gated sodium channel. Brain 138: 891-901

Lu J, McKinsey TA, Zhang CL, Olson EN (2000) Regulation of skeletal myogenesis by association of the MEF2 transcription factor with class II histone deacetylases. Mol Cell 6: 233-44

Ludlam WH, Taylor MH, Tanner KG, Denu JM, Goodman RH, Smolik SM (2002) The acetyltransferase activity of CBP is required for wingless activation and H4 acetylation in Drosophila melanogaster. Mol Cell Biol 22: 3832-41

Mao Z, Bonni A, Xia F, Nadal-Vicens M, Greenberg ME (1999) Neuronal activity-dependent cell survival mediated by transcription factor MEF2. Science 286: 785-90

Marinescu VD, Kohane IS, Riva A (2005) MAPPER: a search engine for the computational identification of putative transcription factor binding sites in multiple genomes. BMC Bioinformatics 6: 79

Markstein M, Pitsouli C, Villalta C, Celniker SE, Perrimon N (2008) Exploiting position effects and the gypsy retrovirus insulator to engineer precisely expressed transgenes. Nat Genet 40: 476-83

McKinsey TA, Zhang CL, Olson EN (2001) Control of muscle development by dueling HATs and HDACs. Curr Opin Genet Dev 11: 497-504

Mee CJ, Pym EC, Moffat KG, Baines RA (2004) Regulation of neuronal excitability through pumilio-dependent control of a sodium channel gene. J Neurosci 24: 8695-703

Menon KP, Sanyal S, Habara Y, Sanchez R, Wharton RP, Ramaswami M, Zinn K (2004) The translational repressor Pumilio regulates presynaptic morphology and controls postsynaptic accumulation of translation factor eIF-4E. Neuron 44: 663-76

Morin S, Charron F, Robitaille L, Nemer M (2000) GATA-dependent recruitment of MEF2 proteins to target promoters. EMBO J 19: 2046-55

Muraro NI, Weston AJ, Gerber AP, Luschnig S, Moffat KG, Baines RA (2008) Pumilio binds para mRNA and requires Nanos and Brat to regulate sodium current in Drosophila motoneurons. J Neurosci 28: 2099-109

Nguyen HT, Bodmer R, Abmayr SM, McDermott JC, Spoerel NA (1994) D-mef2: a Drosophila mesoderm-specific MADS box-containing gene with a biphasic expression profile during embryogenesis. Proc Natl Acad Sci U S A 91: 7520-4

O’Leary T, van Rossum MC, Wyllie DJ (2010) Homeostasis of intrinsic excitability in hippocampal neurones: dynamics and mechanism of the response to chronic depolarization. J Physiol 588: 157-70

Ogryzko VV, Schiltz RL, Russanova V, Howard BH, Nakatani Y (1996) The transcriptional coactivators p300 and CBP are histone acetyltransferases. Cell 87: 953-9

Rikitake Y, Moran E (1992) DNA-binding properties of the E1A-associated 300- kilodalton protein. Mol Cell Biol 12: 2826-36

Rouillard AD, Gundersen GW, Fernandez NF, Wang Z, Monteiro CD, McDermott MG, Ma’ayan A (2016) The harmonizome: a collection of processed datasets gathered to serve and mine knowledge about genes and proteins. Database (Oxford) 2016

Sartorelli V, Huang J, Hamamori Y, Kedes L (1997) Molecular mechanisms of myogenic coactivation by p300: direct interaction with the activation domain of MyoD and with the MADS box of MEF2C. Mol Cell Biol 17: 1010-26

Siemen H, Colas D, Heller HC, Brustle O, Pera RA (2011) Pumilio-2 function in the mouse nervous system. PLoS One 6: e25932

Sivachenko A, Li Y, Abruzzi KC, Rosbash M (2013) The transcription factor Mef2 links the Drosophila core clock to Fas2, neuronal morphology, and circadian behavior. Neuron 79: 281-92

Stilwell GE, Saraswati S, Littleton JT, Chouinard SW (2006) Development of a Drosophila seizure model for in vivo high-throughput drug screening. Eur J Neurosci 24: 2211-22

Taylor MV, Beatty KE, Hunter HK, Baylies MK (1995) Drosophila MEF2 is regulated by twist and is expressed in both the primordia and differentiated cells of the embryonic somatic, visceral and heart musculature. Mech Dev 50: 29-41

Vessey JP, Schoderboeck L, Gingl E, Luzi E, Riefler J, Di Leva F, Karra D, Thomas S, Kiebler MA, Macchi P (2010) Mammalian Pumilio 2 regulates dendrite morphogenesis and synaptic function. Proc Natl Acad Sci U S A 107: 3222-7

Waltzer L, Bienz M (1998) Drosophila CBP represses the transcription factor TCF to antagonize Wingless signalling. Nature 395: 521-5

Waltzer L, Bienz M (1999) A function of CBP as a transcriptional co-activator during Dpp signalling. EMBO J 18: 1630-41

Wharton RP, Sonoda J, Lee T, Patterson M, Murata Y (1998) The Pumilio RNA-binding domain is also a translational regulator. Mol Cell 1: 863-72

Wreden C, Verrotti AC, Schisa JA, Lieberfarb ME, Strickland S (1997) Nanos and pumilio establish embryonic polarity in Drosophila by promoting posterior deadenylation of hunchback mRNA. Development 124: 3015-23

Wu XL, Huang H, Huang YY, Yuan JX, Zhou X, Chen YM (2015) Reduced Pumilio- 2 expression in patients with temporal lobe epilepsy and in the lithium-pilocarpine induced epilepsy rat model. Epilepsy Behav 50: 31-9

